# High-resolution tracking of unconfined zebrafish behavior reveals stimulatory and anxiolytic effects of psilocybin

**DOI:** 10.1101/2023.04.13.536830

**Authors:** Dotan Braun, Ayelet Rosenberg, Ravid Haruvi, Dorel Malamud, Rani Barbara, Takashi Kawashima

**Author notes:** These authors contributed equally to this work (Dotan Braun, Ayelet Rosenberg).

## Abstract

Serotonergic psychedelics are emerging therapeutics for psychiatric disorders, yet their underlying mechanisms of action in the brain remain largely elusive. Zebrafish have evolutionarily conserved serotonergic circuits and subcortical targets such as the brainstem regions and the cerebellum, providing a promising model for studying the subcortical effects of serotonergic drugs. Here, we developed a wide-field behavioral tracking system for larval zebrafish and investigated the effects of psilocybin, a psychedelic serotonin receptor agonist. Machine learning analyses of precise body kinematics identified latent behavioral states reflecting spontaneous exploration, visually-driven rapid swimming, and irregular swim patterns following stress exposure. Using this method, we identified two main behavioral effects of acute psilocybin treatment: [i] increased rapid swimming in the absence of visual stimuli and [ii] prevention of irregular swim patterns following stress exposure. Together, these effects indicate that psilocybin induces a brain state that is both stimulatory and anxiolytic. These findings pave the way for using larval zebrafish to elucidate subcortical mechanisms underlying the behavioral effects of serotonergic psychedelics.

## Introduction

Mood-related mental disorders cast significant socioeconomic impacts on modern societies, and 5% of adults are estimated to suffer from depression globally^1^. The serotonin theory of depression^2^ emerged soon after the discovery of the serotonergic system in the brain in the 1960s^3^. Since then, the serotonergic system has been a therapeutic target for major depression, obsessive-compulsive disorders, and other psychiatric disorders^4^. Current medication regimens based on serotonin-selective reuptake inhibitors (SSRIs) have limited efficacies in terms of quickness of therapeutic effects^5^ and final remission rates^6^, calling for a better understanding of neural mechanisms for mood-related behavioral alternation and its pharmaceutical rescue.

The recent resurgence of the use of hallucinogenic drugs as fast-acting antidepressants has opened new opportunities for the research of neural circuit mechanisms critical for the treatment of mood-related disorders^7^. Psilocybin, a psychedelic compound, originates in the genus of gilled mushrooms *Psilocybe* and acts as a potent agonist for a family of serotonin receptors. Psilocybin is effective for clinical cases of treatment-resistant depression^8,9^, and only a few doses can have lasting effects on depression symptoms for months or even up to a year^9–15^. These reported therapeutic effects are markedly different from the short-lasting effects of other classes of psychedelics such as ketamine^16^, making psilocybin and its derivatives a promising class of drugs for treating mood-related disorders.

We have a limited understanding of the mechanisms underlying the therapeutic effects of psilocybin. Human brain imaging studies show that psilocybin alters the functional connectivity within the default mode network, including the prefrontal cortex and posterior cingulate cortex^15,17–19^. Microscopic observations of the prefrontal cortex in mice suggest that such changes might occur from the induction of new excitatory synapses^20,21^. There have also been efforts to derive HTR2 agonists that can prevent stress-induced behavioral changes without causing hallucination-like behaviors^21–23^. Very few studies have focused on changes in subcortical structures such as the brainstem and the cerebellum, which are enriched for serotonin receptors. Roles of the cerebellum have been implicated in mood-related disorders^24,25^, and psilocybin affects cerebellar neural activity in humans^26^. In general, neural dynamics in these subcortical structures have been challenging to investigate in mammals.

Larval zebrafish is a model for studying subcortical structures that are evolutionarily conserved across vertebrates. Its small size, optical transparency, and genetic accessibility allow optical recording of neural activity across the whole brain at a single-cell resolution^27–31^. It has a conserved raphe serotonergic system in the hind/midbrain^32–39^ in addition to teleost-specific serotonergic nuclei in the hypothalamus^32,33,40^, that allow detailed investigation of the working principles of serotonergic neurons during behavior. Larval zebrafish has also been used to screen the behavioral effect of stress exposure, antidepressants, and genetic mutations^41–44^. However, few published studies have examined the behavioral effects of psychedelics in larval zebrafish^45–47^, and, to date, there have been no published data on the effects of psilocybin. Such scarcity of behavioral insights impedes further research into their actions in neural circuit dynamics.

Tracking of millisecond-timescale body kinematics identified diverse types of latent behavioral states in larval zebrafish^48,49^, and these findings provided direct insights into underlying neural mechanisms^35,50^. Yet, previous studies examining the effect of drug treatments on behavioral stress responses in zebrafish have generally focused on macroscopic parameters such as overall travel distance and environmental preference. The lack of body kinematics information in these studies makes it challenging to connect observed behavioral changes and underlying neural mechanisms. A barrier to using high-speed behavioral tracking to study stress responses is that the small chambers typically used for such behavioral tracking can themselves incur confinement stress^41^. Thus, precision approaches have yet to be explored for studying the effects of drug treatments on stress responses in zebrafish.

To overcome this challenge, we developed a machine-learning approach that tracks body kinematics in a large, unconfining environment and infers changes in behavioral states by stress exposure and psilocybin treatments. This approach enabled us to disambiguate distinct behavioral states governing spontaneous exploration, visually-driven rapid swimming, and irregular swim patterns after stress exposure. Acute psilocybin treatment facilitated rapid swimming in the absence of visual stimuli (stimulatory effect) and prevented occurrences of irregular swim patterns following stress exposure (anxiolytic effect). These behavioral effects parallel clinical observations and open new opportunities for studying how serotonergic psychedelics impact neural dynamics in subcortical structures.

## Results

### High-resolution fish tracking system for a large environment

We developed an experimental setup and a data processing pipeline that examine how innate behaviors of larval zebrafish, such as spontaneous exploration and optomotor response, change after drug treatments and stress exposure. This setup tracks the precise body kinematics of a single fish in an environment that is large (90 mm) compared to the length of larval zebrafish (∼4 mm) at a high-resolution (> 1100 x 1100 pixels) and high-speed (290 Hz) (**Figures 1A and S1A**). Fish behavior was recorded at infrared wavelengths for 15 minutes while visual gratings stopped for 10 seconds (spontaneous swimming) and moved for 10 seconds (visually-driven behavior) in cycles (**Figures S1A and S1B**). We tested a single fish per experiment to exclude the effect of social dynamics throughout this study. The large size of the imaged arena resulted in lower pixel resolution relative to previous body kinematics studies^48–50^. Therefore, our data processing tracked head trajectories and tail kinematics at subpixel resolution, i.e., at spatial scales smaller than the pixel size enabled by prior knowledge of the shape of the animal. We achieved localization accuracy at around 25 μm for the head trajectory (**Figure S1C**).

**Figure 1:**
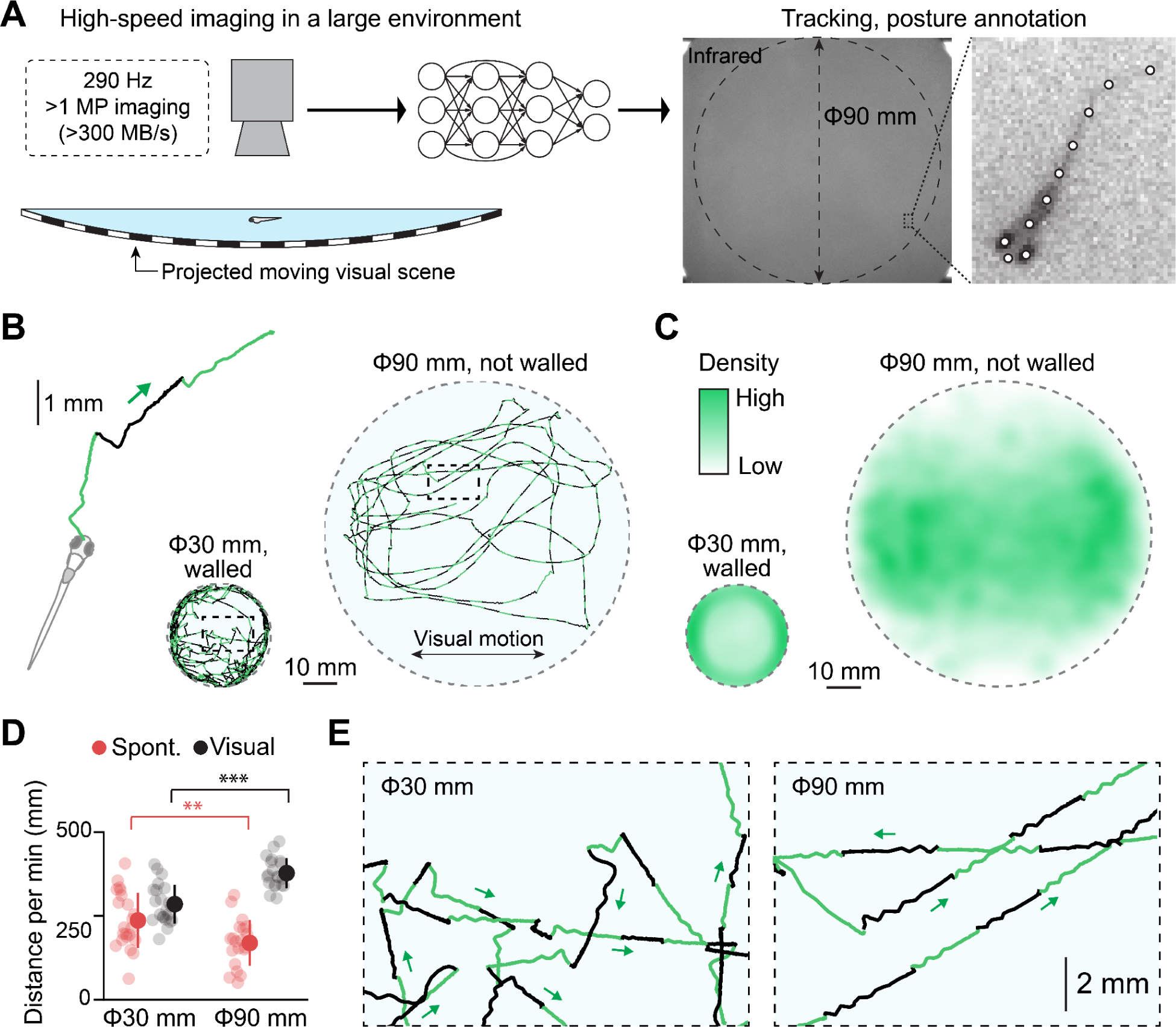
High-resolution, high-speed tracking of zebrafish behavior in a large environment. **(A)** Schematic illustration of our experimental setup and analysis pipeline. Fish swam in an arena where visual stimuli were projected underneath. We acquired high-resolution images at a speed of 290 Hz across the arena of up to 90 mm in diameter. Data analysis pipelines processed the images to identify the fish loci and body postures by using a deep neural network. See Figure S1A and the Methods section for details. **(B)** Head centroid trajectories in a small, walled environment (30 mm) and a large, unwalled environment (90 mm). Each swim episode is colored in either light green or black to visually separate different swim episodes. **(C)** Distributions of head centroids during experiments across tested fish. N=22 and N=20 fish for the small and large dishes, respectively. **(D)** Swimming distance per minute during spontaneous exploration (red) and visually-induced swimming (black). Small circles represent individual fish. **, p=0.0081; ***, p=1.7*10^-6^ from 2-sample t-test. **(E)** Expanded head centroid trajectories from the outlined central parts of the small arena (left) and the large arena (right) from (B). The large environment facilitates straight swim patterns.

Using this setup, we examined behavioral impacts of the size of behavioral arenas by comparing swimming trajectories between those in a small, walled arena (30 mm) and those in a large unwalled arena (90 mm). Zebrafish typically swam near the wall in the small arena due to their innate preferences called thigmotaxis^51^. In our large arena, on the contrary, they explored widely (**Figures 1B and 1C**) and swam longer distances during visual stimulus motion (**Figure 1D**). Spontaneous swimming distance was longer in the small arena, indicating a stimulatory effect of confinement stress^41^ (**Figure 1D**). We also observed notable differences in swim trajectories around the central area of the dish. Fish showed frequent turning in the small arena even when they were not near the wall, potentially due to confinement stress, whereas they swam mostly in straight lines in the large arena (**Figures 1F and S1D**). Fish also showed straight swim patterns in a different large arena with a boundary wall (**Figure S1E**). These observations indicate that the size of the behavioral arena significantly impacts the swim patterns of larval zebrafish.

### Large environment expands repertoires of swim patterns with less confinement artifacts

We quantified tail motions using a deep neural network (EfficientNet-B6^52^) implemented in DeepLabCut software^53^ (**Figure 2A**). We trained the network over 550 images (**Figure S2A**) manually annotated for ten body parts, including the eyes, nostril, body trunk, and six points along the tail. We fitted quadratic curves to the identified points along the tail to quantify tail motions (**Figure S2B**). Lateral movements of the head centroid always synchronized with the tail movements (**Figure S2C**) and were used as a reference for extracting tail motion parameters (**Figure 2B**). We validated the accuracy of quantified tail movements by examining how well tail parameters can predict the swim distance for each swim event. Our prediction model based on extracted tail parameters yielded a Pearson correlation coefficient of 0.89 ± 0.036 across 20 fish, indicating a highly accurate extraction of tail motion parameters (**Figure S2D**).

**Figure 2:**
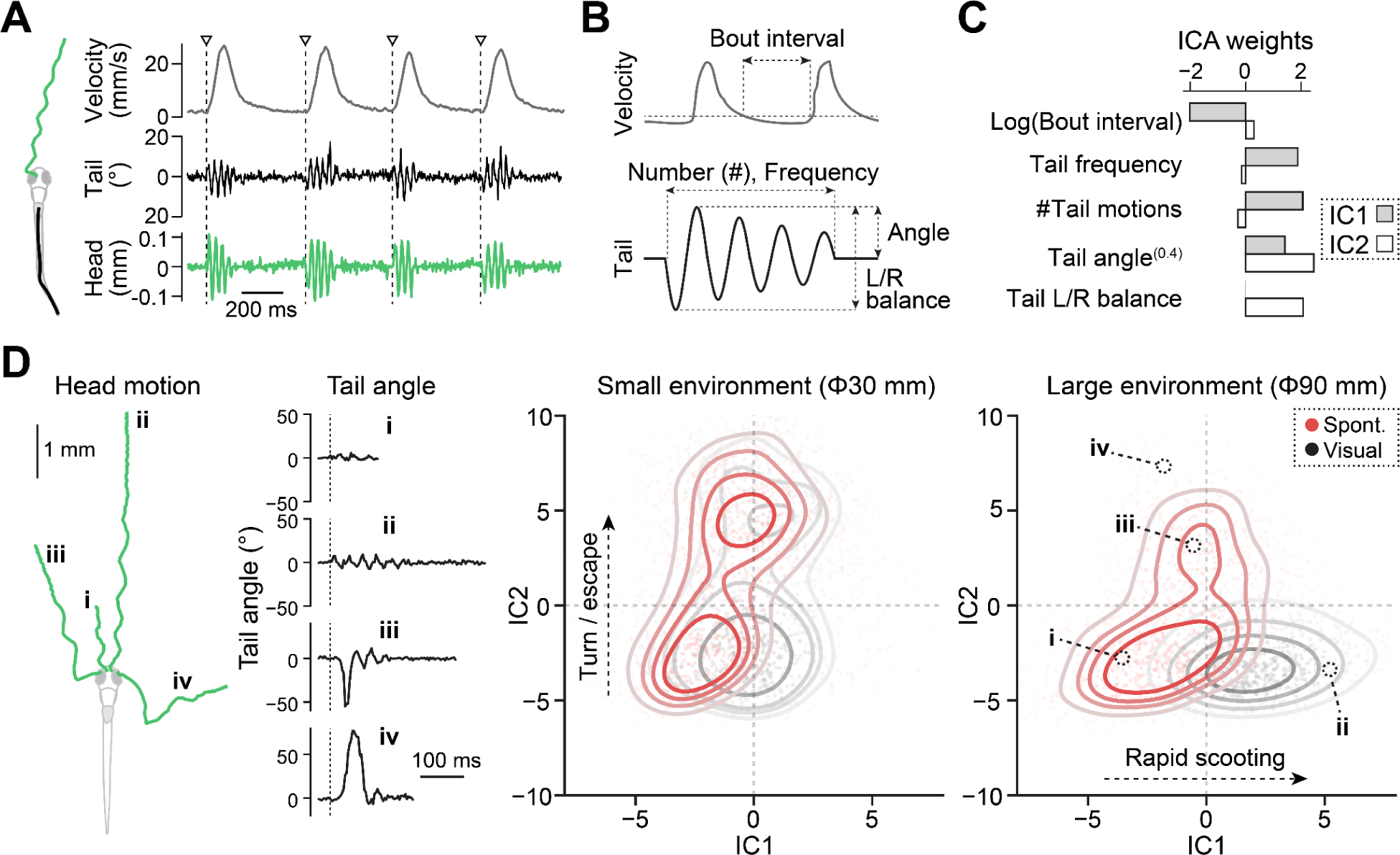
Large environment expands behavioral repertoires with less confinement artifacts. **(A)** *S*wim velocities (gray), tail motion (black) and head centroid motions (green) during multiple swim episodes. Triangles represent the onset of individual swim episodes. **(B)** Swim parameters and tail motion parameters for independent component analyses (ICA). **(C)** ICA weights for component #1 (gray) and component #2 (white). We used tail angle as a power of 0.4 because it best correlated with the swim distance (Figure S2D). Swim parameters and tail motion parameters were Z-scored individually before performing ICA. **(D)** Scatter density plot of various swim patterns in the ICA space reveals enriched repertoires of swimming in a large environment. *Left*, head centroid trajectories and tail motions of four representative swim patterns. *Right*, scatter density plots of swim patterns in the small and large environment during spontaneous exploration (red) and visually-induced swimming (black) in the ICA space. The same number (1200) of randomly selected swim events from N=22 and N=20 fish for the small and large dishes, respectively, were plotted for each condition. Color saturations of dots and contour lines represent the local densities of all the collected data points. The loci of four representative swim patterns on the left are marked in black circles. Larger environment (90 mm) facilitated rapid long scooting (IC1) during visually-driven swimming and fewer turnings/escapes (IC2) during both spontaneous and visually-driven swimming.

We next examined how arena sizes affect latent behavioral states by using independent component analysis (ICA) of five separate parameters of swimming (**Figure 2C**): frequencies of tail motions, angles of tail motions, the number of tail motions, the balance of tail motions between left and right sides, and intervals between swim episodes. Such dimensionality reduction analysis based on multiple parameters yields more robust estimates of latent behavioral states^54^. We applied this analysis to swim episodes in the central part of small and large arenas (**Figure 1**, see Methods) and identified the first two independent components (IC1, IC2) in an unbiased manner (**Figure 2C**). These components allowed us to map various types of swim patterns, including short scooting (i), rapid long scooting (ii), routine turns (iii), and C-turns (iv), into different loci on a low-dimensional space (**Figure 2D**). IC1 separated short scooting during spontaneous exploration and long rapid scooting during visual stimulus motion. IC2 separated scooting and turning/escape behaviors.

This mapping method showed a clear separation of swim patterns during visually-driven swimming from those during spontaneous exploration in the large arena (**Figure 2D**). Such separation was obscure in the small arena. Long rapid scooting (IC1) was dominant during visual stimulus motion in the large arena, whereas turning and escape behaviors (IC2) were more dominant in the small arena (**Figures 2D and S2E**). At the individual parameter level, the large arena allowed higher tail frequency and more tail motions per bout during visual stimulus motion and longer bout intervals during spontaneous exploration (**Figure S2F**), consistent with the above observation of swim trajectories (**Figure 1**). These body kinematics analyses demonstrate that a large arena, which is more than 20 times the body length of larval zebrafish, is essential for evaluating the full extent of swimming repertoires while minimizing confinement-induced turning/escape behaviors.

### Psilocybin has stimulatory effects on spontaneous exploration

We tested the effects of psilocybin treatment on spontaneous exploration and visually-driven swimming by using the above machine learning methods. Psilocybin and its metabolite psilocin act as agonists for serotonin receptors. Upon ingestion, psilocybin converts to psilocin by the action of endogenous phosphatases. Psilocin crosses the blood-brain barrier^55^ and has stronger affinities to serotonin receptors (**Figure 3A**). Psilocin has affinities to multiple types of serotonin receptors, including inhibitory HTR1 receptors and excitatory HTR2 receptors^56–58^. Serotonin receptors in zebrafish are highly similar to those in humans and include major types from HTR1 to HTR7 and their subtypes. Unbiased homology analysis of protein sequences showed robust co-clustering of human and zebrafish serotonin receptors down to subtypes such as HTR2A, 2B, and 2C (**Figures 3B and S3A**). We also confirmed the high expression of zebrafish HTR2 receptors in the brain (**Figure 3C**). Therefore, it is reasonable to hypothesize that psilocybin and its metabolite psilocin have behavioral effects on larval zebrafish.

**Figure 3:**
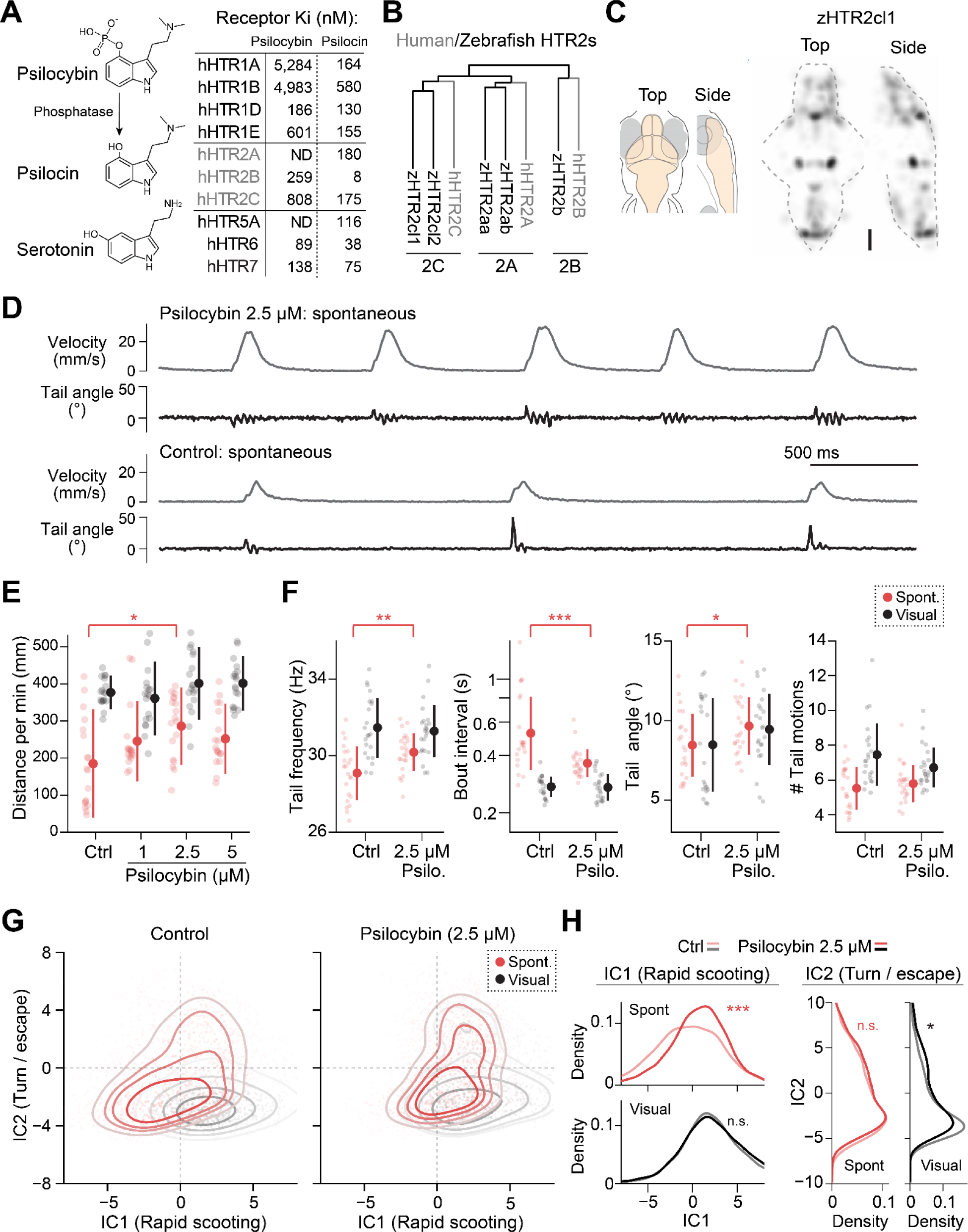
Psilocybin has stimulatory effects on spontaneous exploration. **(A)** Affinities of psilocybin and its metabolite psilocin to human serotonin receptors. Upon ingestion, psilocybin is metabolized by endogenous phosphatases into psilocin, which is structurally similar to serotonin. Psilocin has nanomolar affinities to a wide range of serotonin receptors. Affinities values are taken from a reference^56^. **(B)** *Left*, unbiased homology analysis of protein sequences revealed conserved subclasses of type 2 serotonin receptors between zebrafish and humans. **(C)** Average expression map of HTR2cl1 gene across 5 zebrafish brains obtained by using RNA fluorescence in situ hybridization. Scale bar, 100 μm. See Methods for details. **(D)** Psilocybin evokes rapid scooting behaviors during spontaneous exploration. **(E)** At a concentration of 2.5 μM, psilocybin significantly enhances swimming distances during spontaneous exploration but not visually-driven swimming. N=18 fish for each condition. *, p=0.011 from Tukey’s post-hoc test after one-way ANOVA detected a significant difference (F=3.6) among groups for spontaneous swimming distance. **(F)** Psilocybin significantly enhanced tail frequency, shortened bout intervals, and slightly enhanced tail angles during spontaneous swimming. N=22 and 23 fish for control and 2.5 μM conditions, respectively. P values are from a 2-sample t-test between groups. **, p=0.0051 (frequency); ***, p=8.6*10^-4^ (interval); *, p=0.044 (angle). **(G)** Independent component analysis (ICA) reveals the shift of spontaneous swim patterns toward the distribution of visually-driven swim patterns. The same number (1200) of randomly selected swim events were plotted for each condition. Color saturations of dots and contour lines represent the local densities of all the collected data points. **(H)** Psilocybin significantly enhances rapid scooting (IC1) during spontaneous exploration, while it does not cause a significant increase in turning/escape behaviors (IC2). Statistical analyses of IC1 were performed using the same set of fish in Figure 1 by using kernel density 2-sample test (see Methods). We included 6,128 (control) and 9,395 (2.5 μM) swim episodes for the statistics of spontaneous exploration and 6,413 (control) and 6,427 (2.5 μM) swim episodes for the statistics of visually-driven behavior. ***, p=3.3*10^-65^ and *, p=0.013 between the control group and those after exposure to 2.5 μM psilocybin. n.s., not significant (p>0.05). Error bars represent standard deviations across tested fish.

We found that acute, short bath pretreatment with psilocybin (2.5 μM, 4h) in larval zebrafish had stimulatory effects on spontaneous exploration. We determined this pretreatment protocol after testing dosages between 1 uM and 50 uM and durations between 30 minutes to 24 hours (**Figure S3B**). The concentration of 2.5 uM amounts to a slightly higher dosage (0.71 mg/kg) compared to the clinical dosage in humans (0.6 mg/kg)^59^. This optimal duration of 4h is consistent with the time course of passive diffusion of a drug with similar molecular weight into the brain of larval zebrafish^60^. We observed the reduction of such effects at higher concentrations and longer durations, indicating that the action of psilocybin becomes saturated at this relatively low concentration compared to serotonin-selective reuptake inhibitors (see below). This saturation effect is consistent with clinical observations in human subjects^61^.

After pretreatment with psilocybin, fish swam with shorter intervals with faster velocities (**Figure 3D**), resulting in enhanced swim distance during spontaneous exploration (**Figure 3E**). They showed significantly enhanced tail frequencies and shorter intervals between swim bouts similar to those observed during visual stimulus motion (**Figure 3F**). We did not see noticeable changes in these parameters during visual stimulus motion, indicating that psilocybin’s effect is limited to spontaneous exploration in our paradigm. Independent component analysis of swim parameters also confirmed the above observation (**Figure 3G**). We observed a significant shift in spontaneous swim patterns toward the direction of visually driven rapid scooting along the axis of IC1 (**Figure 3H**). These results indicate that psilocybin stimulates swim patterns in a partially similar manner to visual stimuli.

The effect of acute exposure to psilocybin was different from that of serotonin-selective reuptake inhibitors (SSRIs) that block serotonin reuptake and increase serotonin concentration at synapses^62^. Full therapeutic effects of SSRIs do not occur during acute dosage in humans^5^, and some studies showed elevated anxiety levels during the first few weeks of SSRI treatment^63^. Consistently with previous reports in zebrafish^64–66^, we observed that pretreatments with fluoxetine and fluvoxamine have suppressive effects on swim patterns (**Figure S3**). While fluoxetine caused noticeable distortions in the swim patterns and low-frequency tail motions, fluvoxamine caused decreases in the amplitudes of tail motions (**Figure S3C**). Swimming distances decreased linearly with the dosage for both spontaneous exploration and visually-driven swimming, indicating that SSRIs suppress motor circuits in the brain regardless of external stimuli (**Figure S3D**). These differences in behavioral effects between psilocybin and SSRIs suggest that psilocybin’s stimulatory effects may occur from its selective affinities to a subset of serotonin receptors (**Figure 3A**).

### Psilocybin prevents stress-induced changes in swim patterns

Acute administration of psilocybin has anxiolytic effects in humans^11,67^. We thus tested whether acute stress exposure causes changes in fish’s swim patterns in our setup and whether psilocybin can prevent such stress-induced behavioral changes. Various environmental stressors have been tested in larval zebrafish that trigger cortisol increase, including hypertonic water^43,68–74^, acids^68^, mechanical disturbance^43,70,74,75^, social isolation^70^, and heating^70,76^ / cooling^70^ shock. Here we used an acute cold shock protocol^70^ that rapidly lowers the temperature by 10 degrees (28**°**C to 18**°**C, **Figure 4A**) as it is least likely to cause physiological stress from lasting changes in tissue integrities, protein folding, and ionic balance in the body^70^. We also tested the effect of hypertonic stress for comparison (**Figure S4**). We pre-treated fish with psilocybin with the most effective concentration for enhancing spontaneous exploration (2.5 μM, **Figure 3**), exposed them to stressors for 5 minutes, recovered them at a normal temperature, and tested their spontaneous exploration and visually-driven swimming (**Figure 4A**). Exposure to both cold and hypertonic stressors increased the swimming distance (**Figures 4B and S4A**), which is consistent with the previous reports^68,69^ and demonstrates the robustness of our stress protocol.

**Figure 4:**
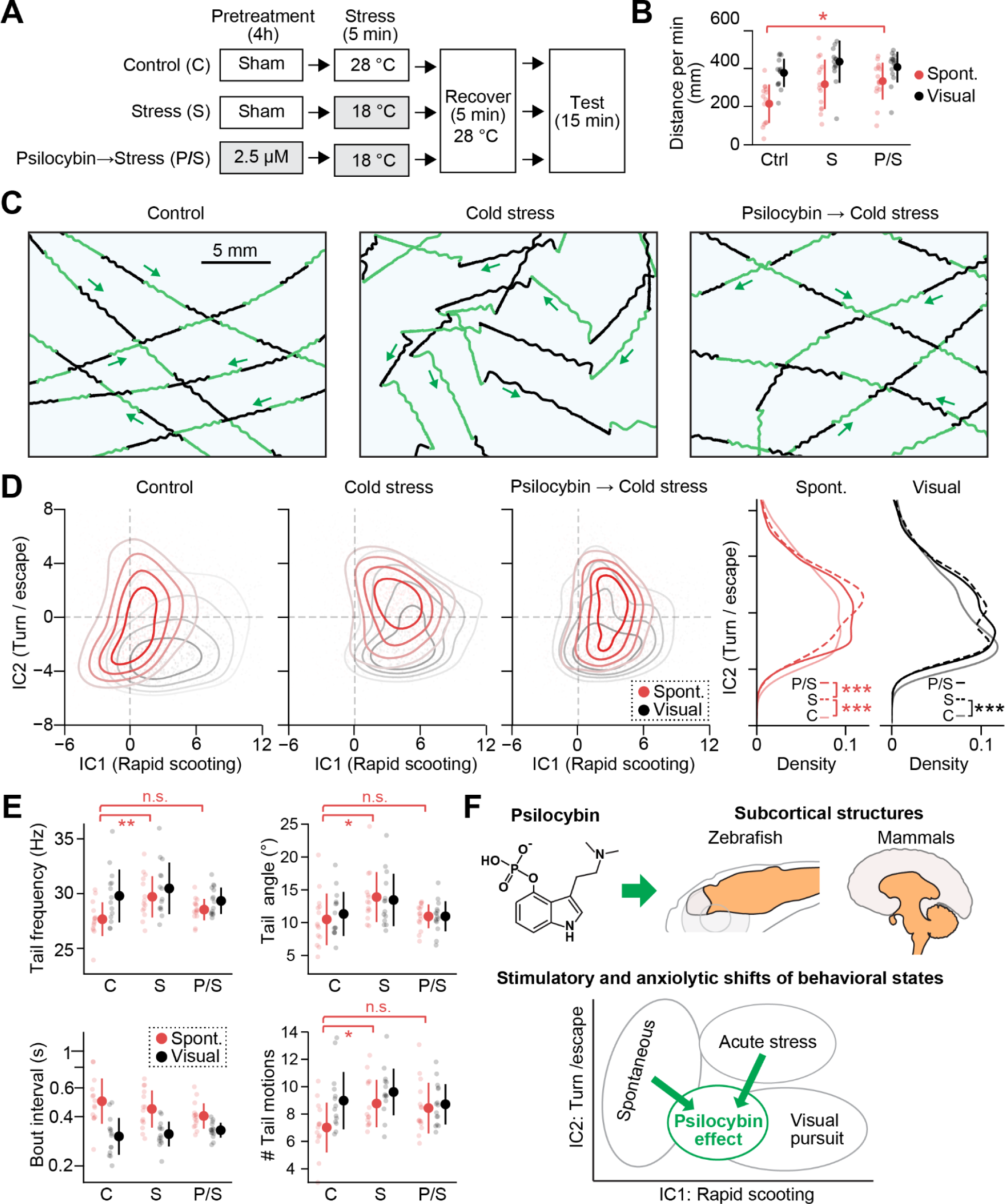
Psilocybin prevents stress-induced behavioral changes. **(A)** Behavioral paradigms for psilocybin treatment and acute cold stress exposure. **(B)** Swimming distances during spontaneous exploration (red) and visually driven swimming (black). N=14 (C), 14 (S), 16 (P) and 15 (P/S) fish. *, p=0.021 from Tukey’s post-hoc test between the control and P/S conditions after one-way ANOVA detected significant differences among groups for spontaneous swimming distance. **(C)** Acute cold stress exposure induced zigzag swim patterns during visually-driven swimming, and psilocybin prevented such stress-induced behavioral changes. Shaded boxes indicate changed temperature or drug conditions. **(D)** *Left*, independent component analysis (ICA) revealed the shift of swim patterns toward turn/escape behavior (IC2) after cold stress. Pretreatment with psilocybin prevented such a shift. The same number (1200) of randomly selected swim events were plotted for each condition. Color saturations of dots and contour lines represent the local densities of all the collected data points. *Right,* statistical analyses of the occurrences of turning/escape behaviors based on the IC2 component. We used kernel density 2-sample test (see Methods). We included 4,216 (C), 3,833 (S) and 4,872 (P/S) swim episodes for the statistics of spontaneous exploration, and 3,312 (C), 2,713 (S) and 2,706 (P/S) swim episodes for the statistics of visually-driven behavior. ***, p=1.8*10^-31^ (C vs S) and 3.0*10^-8^ (S vs P/S) during spontaneous exploration. ***, p=1.0*10^-7^ (C vs S) during visually driven swimming. **(E)** Analyses of individual swim parameters. Acute cold stress exposure significantly increased tail frequency, tail angle and the number of tail motions, while such an effect was not observed in fish pretreated with psilocybin. Statistical tests used Tukey’s post-hoc test after one-way ANOVA among different conditions during spontaneous explorations: **, p=0.0035 (tail frequency); *, p=0.032 (tail angle); *, p=0.046 (tail motions). Error bars represent standard deviations across tested fish. Error bars represent standard deviations across tested fish. **(F)** Summary of our findings in this study. Psilocybin may act on subcortical structures that are evolutionarily conserved between fish and mammals. It stimulates spontaneous behaviors and prevents stress-induced behavioral changes by creating an intermediate behavioral state between spontaneous exploration and visually-driven swimming in larval zebrafish. The human brain is a modification of a drawing by Patrick J. Lynch.

We found that acute stress exposure caused “zigzag” swimming patterns compared to the control (**Figure 4C**). During visual stimulus motion, the control fish shows straight trajectories in our large arena (**Figure 1F**). Such patterns changed into zigzag patterns, as each bout started from a sharp turning of the head and large tail undulation to one side (pattern [iii] in Figure 2D) instead of smooth scooting (patterns [i] or [ii] in Figure 2D). This type of change was also observed after exposure to hypertonic stress (**Figure S4B**), suggesting that the emergence of zig-zag swim patterns, as compared to normal smooth straight patterns, can be a robust indicator of stress-induced behavioral changes in larval zebrafish.

Importantly, pre-treatment with psilocybin prevented stress-induced changes in swim patterns. Psilocybin-pretreated fish exhibited straight swim patterns even after the stress exposure (**Figure 4C**). This prevention of stress-induced behavioral changes was also evident in the ICA analysis along the axis of IC2, which represents occurrences of escape/turning behavior (**Figure 4D**). Acute cold stress significantly elevated distributions along the IC2 axis (**Figure 4D**). The pretreatment with psilocybin significantly diminished the increased occurrence of turning/escape behavior after cold stress (**Figure 4D**). Such preventative effect was also evident in individual tail kinematics, such as frequencies and angles of tail motions (**Figure 4E**). Notably, psilocybin pretreatment did not prevent the stress-induced shift of behavioral states along the IC1 axis, which represents occurrences of rapid scooting behavior (**Figure 4D**). This shift along the IC1 axis is consistent with the effect of psilocybin pretreatment per se (**Figure 3G**). These results suggest that the stimulatory effect of psilocybin prevents the stress-induced occurrence of escape/turn behavior at the same time.

We did not observe similar preventative effects of psilocybin for behavioral changes induced by hypertonic stress (**Figure S4D**). This discrepancy is potentially due to the difference between central, anxiety-like stress and physiological stress for larval zebrafish^70^ and indicates that psilocybin’s anxiolytic effects may result from preventing the occurrence of anxiety-related neural dynamics rather than from inducing straight swim patterns at the motor circuit level.

We further examined the involvement of HTR2 receptors in stress-induced behavioral changes by using ketanserin, an HTR2 receptor antagonist^77^ which also inhibits monoamine transporters^78^ and histamine receptors^79^. Ketanserin has acute anxiogenic effects in adult zebrafish^80,81^ and rodents^82,83^. We found that bath application of ketanserin shifted behavioral states similarly to both cold and hypertonic stressors, and psilocybin prevented such changes (**Figure S4E**). These results indicate the crucial roles of HTR2 receptor pathways in behavioral changes following stress exposure and demonstrate that the observed stimulatory and anxiolytic effects of psilocybin likely occur from the modulation of such endogenous serotonergic pathways in the zebrafish brain.

## Discussion

In this study, we developed a high-resolution tracking system and a machine-learning framework for evaluating how psilocybin changes the latent behavioral states of larval zebrafish. Psilocybin has stimulatory effects on spontaneous swimming and preventative effects for stress-induced behavioral changes. These effects converged toward an intermediate state between spontaneous exploration and visually-driven rapid swimming in the latent behavioral space, indicating that psilocybin induces unique neural dynamics that are both stimulatory and anxiolytic (**Figure 4F**). These observations have similarities with those in mammals, indicating the presence of common neural mechanisms in the evolutionarily conserved brain areas through which psilocybin exerts its behavioral effects.

Tracking precise body kinematics in a larger environment was critical for our findings in this study. Our 90-mm arena facilitated straight swim patterns compared to frequent turning/escape behaviors in a small arena, and it allowed us to identify distinct behavioral states that affect spontaneous exploration, visually-driven rapid scooting and irregular swim patterns after stress exposure. These results indicate that high-throughput assays in small arenas such as multiwell plates may impair the full spectrum of the fish’s behavioral repertoires. Moreover, we demonstrated that environmental stimuli that evoke different types of body kinematics, such as moving stimuli and acute stress exposure, could yield similar changes in macroscopic locomotion measures such as swim distance per minute (**Figures 1, 4 and S4**). Precise tracking of body kinematics enabled robust inference of the shifts of latent behavioral states during stress exposure and psilocybin administration.

Our observations open up new opportunities for further investigations into subcortical neural mechanisms by which psilocybin affects behaviors. While psilocybin is known primarily as an agonist for type 2 serotonin receptors, it also activates HTR1 receptors to exert its behavioral effects^57,58^. Type 1 receptors are densely expressed in the brainstem areas of mammals^84,85^ and zebrafish^86^, whereas type 2 receptors densely express in cerebellar areas of mammals^87–89^ and zebrafish^90^. Therefore psilocybin likely acts on these receptors to alter neural dynamics in the brain of zebrafish.

How does psilocybin stimulate swimming in a partially similar manner to visual stimuli? Neural mechanisms that trigger spontaneous swimming remain mostly elusive in zebrafish. Recent studies found that spontaneous activation of a sensory neural ensemble in the optic tectum triggers spontaneous swimming^91^. Therefore, it is possible that psilocybin stimulates a part of the sensorimotor reflex circuit to induce swim patterns that are partially similar to those during visual stimulus motion (**Figure 3**). It is also possible that persistent activation/suppression of motor circuits underlies such behavioral changes. Optogenetic activation of cerebellar Purkinje neurons induced swimming in zebrafish^92^ and locomotion in mice^93^. Further investigation into neural dynamics based on whole-brain neural activity imaging methods^27,36,94^ and histological neural activity mapping methods^40^ will be necessary to disambiguate these potential mechanisms.

Psilocybin’s preventative effects for stress-induced behavioral changes may occur from the same neural mechanisms responsible for its effect on spontaneous explorations, as we found that both seem to bring the fish’s swim patterns toward an intermediate state between spontaneous exploration and visually-driven rapid scooting (**Figure 4F**). Such common mechanisms can occur at neural circuit levels and molecular levels^95^. Acute administration of psilocybin increases cortisol levels in humans despite its anxiolytic effects^67,96^, indicating that psilocybin may not act directly on the hypothalamic-pituitary-adrenal axis and rather makes neural dynamics in the brain resilient to stress exposure. Such effects may occur through the cerebellum system. Patients with cerebellar neurodegeneration suffer from depressive symptoms^97^, and the therapeutic effects of cerebellar stimulation have been clinically demonstrated^24,25,98^. Chronic administration of serotonin-reuptake inhibitors increased functional connectivity between the cerebellum and midbrain structures in depression patients^99^. These insights point to the pivotal roles of serotonergic modulation of the cerebellum in mood-related disorders. To address these questions, further investigations into brain-wide neural dynamics during acute stress exposure and psilocybin treatment are necessary, which can be done by implementing these behavioral paradigms into whole-brain imaging setups for head-fixed zebrafish^28,36,68,100^ or freely swimming zebrafish^29,101^.

This study only focused on the acute effect of psilocybin in larval zebrafish and did not address other clinically relevant actions of psilocybin. For example, psilocybin and other HTR2 agonists are effective in reversing depression-like behaviors after chronic stress exposure in rodents^21–23^ and humans^9–15^, and a few doses have lasting effects^9^. The latter persistent effect is unique to psilocybin compared to other antidepressants such as ketamine and SSRIs, but its underlying mechanisms are largely unknown. Our findings pave the way to examine serotonergic psychedelics’ unique pharmacological actions in the brains of larval zebrafish, which allow for live tracking of neural activity^27,35,36^, neurotransmitters^102,103^, structural plasticity^104^, and molecular dynamics^105^ across the brain.

## Methods

### Animal experiments

All of the experiments in this study that use zebrafish larvae were performed under the approval of the Institutional Animal Care and Use Committee (IACUC) and the Institutional Biosafety Committee (IBC) of the Weizmann Institute of Science and by the Israeli National Law for the Protection of Animals - Experiments with Animals (1994). The use of psilocybin in our research was conducted under the supervision of the Israel Ministry of Health (number # 984.02.2022) to D.B. and the IACUC (number # 07550922-2) of the Weizmann Institute of Science.

### The hardware of the zebrafish tracking system

We used a custom-built zebrafish tracking system consisting of a high-speed camera (FLIR, ORX-10G-51S5M-C monochromatic), a macro lens and its locking sleeve (ZOOM 7000 & 1-11736, Navitor), an infrared filter for the lens (Hoya R-72, Edmund #54-753), 880 nm LED illumination(100 x 200 mm, BL040801-880-IC, Advanced Illumination; Edmund # 66-844), a power supply and manual intensity adjustments for the LED (Edmund # 86-887, #66-855), a 100 mm x 120 mm cold mirror (Edmund #64-452), and a compact projector (Optoma LV130). Structural parts that hold these devices were purchased from Thorlabs or manufactured by the Physics Core Facility of the Weizmann Institute of Science.

The acquisition PC was equipped with a 10G Ethernet adaptor (Intel X540-T2) for data transfer from the camera and an NVMe disk drive that has a 15-terabyte capacity and >3GB/s writing speed (Micron 9300). A custom-written Python application based on PyQT5 (https://pypi.org/project/PyQt5/) and pyqtgraph libraries (https://www.pyqtgraph.org/) controlled devices and acquired data from the camera. Many modules of the application were inherited from PyZebrascope software^94^ which we previously developed for light-sheet microscopy. Communication and data acquisition with the camera used PySpin library (https://pypi.org/project/pyspin/). Raw 8-bit data of the acquired image was saved in a single AVI file for each experiment using OpenCV-Python library (https://pypi.org/project/opencv-python/).

We tested the behavior of *AB* fish at the age of 5 days post fertilization (dpf) on a chemical watch glass (125 mm, AlexRed) whose bottom surface was manually coated with a white spray. We imaged an area that spanned 90 mm (1100-1200 pixels) for each dimension with a resolution of 83 μm per pixel at 290 Hz. We recorded the behavior of each fish (one fish per experiment) for 15 minutes, which resulted in >250,000 frames and ∼ 300 GB file size for each experiment. Behavioral data presented in this study were acquired using at least three batches (**Figures 1, 2, 3, 4, S1D, S2D, S2E, S2F, S3D, S4**) or at least two batches (**Figure S3B**).

### Pharmacological treatments

Until 5 dpf, both control and drug-exposed embryos were reared in 90-mm Petri dishes (Miniplast, 820-090-01-017) and maintained in a light-cycled incubator at 28.0 °C. Media was changed every other day and no food was given before behavioral experiments. All behavioral experiments in this study were performed in E3 medium after washing out pretreated drugs.

#### Psilocybin administration

Psilocybin solution (Sigma, P-097) was purchased as a stock solution concentration of 1.0 mg/mL in acetonitrile:water (1:1), ampule of 1mL (3.52 mM). Stock solutions were stored at −80°C for long-term storage, and in-use aliquots were stored at −20°C. Aliquots were thawed and vortexed immediately before use. Psilocybin was administered by incubating the fish in the psilocybin solution in 6-well plates (Corning, #3516). Desired concentrations of psilocybin were achieved by adding the stock solution to the E3 medium in which the fish swim. Concentrations ranging from 1 μM to 50μM were tested in order to optimize the dosage. We determined that a dose of 2.5 μM for 4 h had the largest effect on spontaneous swimming compared to controls (**Figures 3E and S3B**). Following the 4-hour exposure, fish were double washed in E3 medium, and remained in E3 medium until their behavior was examined. Behaviors were recorded for 15 minutes per fish.

#### SSRI administration

Fluoxetine hydrochloride (Sigma, F918) was prepared as a 1 mg/ml (2.9 mM) stock solution in a conditioned E3 medium for zebrafish embryos. Fluvoxamine (Sigma, F2802) was prepared as a 10 mg/ml (23 mM) stock solution in a conditioned E3 medium. Desired Fluoxetine and Fluvoxamine concentrations were achieved by diluting the stock solution with conditioned E3 medium and storing the aliquots at −20°C. At 5 dpf the fish were incubated as follows: Fluoxetine and Fluvoxamine were administered by incubating the fish in the solutions in 6-well plates. Desired concentrations of the two types of SSRIs were achieved by adding the stock solutions to the E3 medium in which the fish swim. Concentrations ranging from 1 μM to 10μM for Fluoxetine and 2.5μM to 25μM for Fluvoxamine were tested in order to optimize the dosage. After 24 hours of incubation the fish, after being triple washed, were transferred to a new Petri dish, similar to the one they dwelled in before, containing only E3 medium. Behaviors were recorded at 6 dpf for 15 minutes per fish.

#### Stress paradigms

We utilized two established larval zebrafish stress paradigms to induce stress in the fish; hyperosmotic stress and cold temperature stress.

Zebrafish are freshwater fish, and thus a salt-water environment induces psychological and physiological stress^69^. In order to create an osmotic environment, NaCl was dissolved in E3 medium in concentrations of 25 mM, 50 mM, and 100 mM NaCl, which have previously been found to induce stress in zebrafish^43^ (**Figure S4A**). Fish were placed in the osmotic solution for 15 minutes and were then triple-washed immediately before behavioral recordings.

Just as osmotic changes to the water cause stress to the fish, changes in water temperature are also known to induce stress. Zebrafish’s optimal environment is approximately 28°C, and so it has been shown that a short exposure of 5 minutes to 18°C leads to increased cortisol levels and anxiety-like behaviors in larval zebrafish^70^. Thus, we utilized this established stress paradigm and exposed the fish to 18°C E3 medium for 5 minutes. We then returned the fish to 28°C E3 for a 5-minute recovery before testing (**Figure 4A**). 18°C E3 was achieved by mixing 4°C E3 with 28°C E3. Over the course of the 5-minute cold stimulus, the water temperature on average ranged from 18°C to 20°C. The recovery in 28°C E3 was done in an incubator, thus the temperature remained stable. Note that these procedures involve multiple occasions of transferring fish and replacing liquid around the fish, which themselves cause stress response due to mechanical disturbances. The effect of such procedural stress was present as the shifts of behavioral states along the IC2 axis in the data presented in Figures 4 and S4 compared to other datasets.

For ketanserin experiments (**Figure S4E**), we performed a bath application of ketanserin (11.25 μM) for 5 hours before behavioral tests. Psilocybin (2.5 μM) was added to the solution 1 hour after the start of ketanserin treatment for a total duration of 4 hours before behavioral tests. Ketanserin (Sigma, S006) was prepared as a 0.5 mg/mL stock solution in E3 medium. Stock solutions were stored at −80°C for long-term storage, and in-use aliquots were stored at −20°C. The desired concentration of ketanserin was achieved by adding the stock solution to the E3 medium in which the fish swim.

### Tracking zebrafish and its tail motion

We used custom Python scripts to extract swimming parameters and tail movements. We applied the following procedures to each data.

In the first step, we identified the pixel-level centroid position of the head of the fish for each frame and extracted square patches around the fish. To do this, we calculated the average background image based on 100 images equidistantly sampled from all time points and subtracted it from the movie. We applied a gaussian blur filter (σ = 250 μm) to background-subtracted images and identified the darkest pixel as the centroid position of the head of the fish. We cropped the image (144 by 144 pixels, 12 x 12 mm) around the centroid, rescaled it to between 0-255 brightness values, and stored them in a separate AVI file. To accelerate file processing time, we used a recurrent algorithm that searched proximities of the fish positions of previous time points.

In the second step, we automatically annotated body parts from the extracted fish images by using a deep neural network (EfficientNet-B6) implemented in the DeepLabCut package^53^. We trained the network by using 550 manually annotated images (**Figure S2A**). During the development process, we observed that biases in the training datasets were reflected in the automatic annotation results. Therefore we balanced the repertoires of training images so that they covered all angles of the tail symmetrically between the left and right sides. We also included images with outlier pixels, which result from a small inhomogeneity of the bottom coating of the dish. Errors in manual annotations were further screened by independent algorithms that detect significant deviations in distances between annotated body parts and manually corrected. The Deep neural network was trained for 30,000 iterations with “imgaug” augmentation option. The average test error was 1.47 pixels.

In the third step, we applied [i] subpixel head centroid detection and [ii] tail angle quantification for each frame. For subpixel head centroid detection (**Figure S1C**), we applied a gaussian blur filter to the fish image and applied a subpixel centroid detection algorithm implemented in Photutils package (https://photutils.readthedocs.io/) which was originally developed for detecting subpixel motions of cosmic stars in telescopic images. This identification of subpixel-level centroid position was critical for visualizing swim trajectories throughout this study and also for extracting tail motion parameters during swim bouts. For tail angle quantification (**Figure S2B**), we fitted a quadratic function to seven annotated points along the body trunk and the tail and quantified the angle of the fit function relative to the body-nostril axis. The angle quantification was made at 1 mm from the base part of the tail. We tried several other ways of tail angle quantifications from previous studies and found that quadratic fit provides the best signal-to-noise ratio in this low-resolution imaging system.

In the last step, we identified each swim bout based on swim velocities and quantified basic swim parameters (position, velocity, duration, moved distance, etc.) and tail kinematic parameters (frequency, number, angles of tail motions). We found that lateral motions of subpixel head centroid relative to the direction of the swimming, which synchronously precedes tail motions, provide a slightly better signal-to-noise ratio for defining individual cycles of tail motions in this low-resolution imaging system. Therefore, our algorithm used the motion of the subpixel head centroid as a reference to read out maximum tail angles for each tail motion cycle (**Figure S2C**). Peaks of tail angles are detected for each tail motion cycle and further averaged across cycles to quantify average tail angles. Tail frequencies are quantified based on these tail motion cycles. The left / right (L/R) balance of the tail motion is calculated by using the below equation

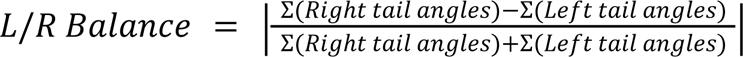

so that the value becomes 0 if the tail motions are symmetric and 1 if there is only one tail movement to a specific side. We tested the accuracy of tail motion tracking by examining how accurately we can predict the swim distances from tail motion parameters (**Figure S2D**). We created a multiplicative model that predicts the distance of swimming based on the frequency, number, and average angle of tail motions for each swim bout. We extracted forward swim events that occurred in the central part of the large dish (30 mm from the center) and caused changes in the head direction less than 30 degrees. We fit the model by using the L-BFGS-B method implemented in the minimize function of Scipy package and identified optimal values for the power factors for each tail parameter across 20 fish. The optimal power factors were roughly 0.4 for the tail angle and 1 for the frequency and the number of tail motions. We then calculated the correlation coefficient with the actual swim distance and the predicted distance from tail motions (**Figure S2D**).

### Data analysis of tail motions

We used custom Python scripts to summarize the results presented in this study. Except for the analysis of swimming distances (**Figures 1D, 3E, 4B, S3B, S3D and S4A**), we only included swim episodes that occurred in the central part of behavioral arenas (within 10 mm or 30 mm from the center of the small or large arena, respectively) for the analyses of swim/tail parameters throughout this study to rule out the physical effect of the wall in the small arena and shallow places in the large arena.

Statistical tests for swimming distances, head angle changes and individual tail parameters (**Figures 3F, 4E, S1D, S2F, S3D**) used either 2-sample t-test or Tukey’s post-hoc test followed by one-way analysis of variance (ANOVA) in the Scipy package (https://scipy.org/).

For independent component analysis (ICA) (**Figures 2D, 3G, 4D, S4C and S4D**), we first identified ICA weights and normalization factors in the dataset of 14,694 swimming episodes from N=22 fish for the small arena and N=20 fish for the large arena (**Figure 1**) by using FastICA function in scikit-learn package (https://scikit-learn.org). Then we applied the same ICA weights and normalization factors to other datasets. Scatter density plots in this study (**Figures 2D, 3G, 4D, S4C, S4D and S4E**) were generated using *gaussian_kde* function in Scipy package for dot coloring and *kdeplot* function in Seaborn package (https://seaborn.pydata.org/) for contour lines. Statistical tests for distributional differences of independent components (**Figures 3H, 4D, S2E, S4C, S4D, S4E**) were performed using kernel density 2-sample test (kde.test) in R package through rpy2 Python-R bridge (https://pypi.org/project/rpy2/). We used this density-based test instead of Kolmogorov– Smirnov test because the former provides more conservative levels of significance for larger sample sizes^106^.

### Homology analysis of serotonin receptors

The sequence homology analysis of serotonin receptors between zebrafish and human (**Figures 3B and S3A**) was performed using Clustal Omega^107^ algorithm and iTOL visualization tool^108^ on the EMBL website. Uniplot IDs used for this analysis are as follows. Human serotonin receptors: hHTR1A (P08908), hHTR1B (P28222), hHTR1D (P28221), hHTR1E (P28566), hHTR1F (P30939), hHTR2A (P28223), hHTR2B (P41595), hHTR2C (P28335), hHTR3A (P46098), hHTR3B (O95264), hHTR3C (Q8WXA8), hHTR3D (Q70Z44), hHTR3E (A5X5Y0), hHTR4 (Q13639), hHTR5A (P47898), hHTR6 (P50406), hHTR7 (P34969). Zebrafish serotonin receptors: zHTR1aa (A0A8M1NIJ6), zHTR1ab (A0A8M1NRS3), zHTR1b (B3DK14), zHTR1d (A0A8M2B5P5), zHTR1e (A0A8M9P2V8), zHTR1fa (A0A8M6Z176), zHTR1fb (A0A8M2B6K6), zHTR2aa (A0A8N7TD42), zHTR2ab (A0A8M3B093), zHTR2b (Q0GH74), zHTR2cl1 (A0A8M6Z717), zHTR2cl2 (A0A8M1PZA4), zHTR3a (A0A8M9PD95), zHTR3b (A0A8M9PJB8), zHTR4 (A0A8M9QPE9), zHTR5aa (A0A8M1NJ85), zHTR5ab (Q7ZZ32), zHTR6 (A0A8M3ANX4), zHTR7a (A0A8N7T7N6), zHTR7b(A0A8M9QGY4), zHTR7c (A0A8M1RQY0).

### Histology

zHTR2cl1 expression map (**Figure 3C**) was constructed using RNA fluorescence in situ hybridization (RNA-FISH) with hybridization chain reaction (HCR) method^109^. HCR probe for zHTR2cl1, amplifiers and buffers were purchased from Molecular Instruments. The staining was performed according to an HCR protocol provided by Dr. Inbal Shainer (Max Planck Institute for Biological Intelligence, Germany)^110^ by using a B3 amplifier with Alexa Fluor 546. We used 5-day old transgenic zebrafish that pan-neuronally express nuclear-localized, genetically-encoded calcium indicator (Tg(HuC:H2B-GCaMP7f))^111^ for the staining to register the volumetric image to a reference brain based on GCaMP expression.

Labeled fish were imaged in our custom light-sheet microscope^94^. Prior to the imaging, fish were embedded in 2% low-melting agarose in fish water on a custom-made pedestal inside a glass-walled chamber. Agarose around the head was removed with a microsurgical knife (#10318-14, Fine Science Tools) to minimize the scattering of the excitation laser. For each fish, we acquired two images: Tg(HuC:H2B-GCaMP7f) channel and the zHTR2cl1 HCR staining image.

Imaging data were processed on a Linux server in High Performance Computing (HPC) division at the Weizmann Institute of Science described in our previous work^94^. We performed data processing using custom Python scripts. All the analyses of imaging data were performed on a remote JupyterLab environment (https://jupyterlab.readthedocs.io/). Briefly, we created an average image for each fish. We then identified individual neurons that express nuclear-localized GCaMP based on the average image by using an algorithm for detecting circular shapes in images. Then, we registered the GCaMP image channel of each fish to Tg(elavl3:H2b-GCaMP6s) reference image from mapZebrain database^112^ using Advanced Normalization Tools (ANTs)^113^, and applied the same registration to the HCR in situ image. For each fish, we analyzed the HCR signal in the coordinates of the recognized neurons after subtracting the local background and created a binarized image of the signal. Then, we overlaid a spherical gaussian filter of the positive neurons of each fish, normalized by the number of cells, to create a generalized 3D image of expressing regions.

## Acknowledgment

We thank Inbal Shainer and other members of Kawashima laboratories for experimental help, members of Veterinary Resource at Weizmann Institute of Science for animal care, Yavin Shaham, Marta Moita, Herwig Baier and Ziv Williams for helpful discussion on the project, and Gil Levkowitz, Misha Ahrens, and Sasha Devore for critical reading of the manuscript. This research is supported by Azrieli faculty fellowship (T.K.), Israel Science Foundation Individual Grant (688/22), Binational science foundation (NSF-BSF CNCRS, #2021746), Dan Lebas & Ruth Sonnewend (T.K.), Jonathan Birnbach (T.K.), and Internal grant from the Center for New Scientists at Weizmann Institute of Science (T.K.).

---

**Supplementary Figure 1:**
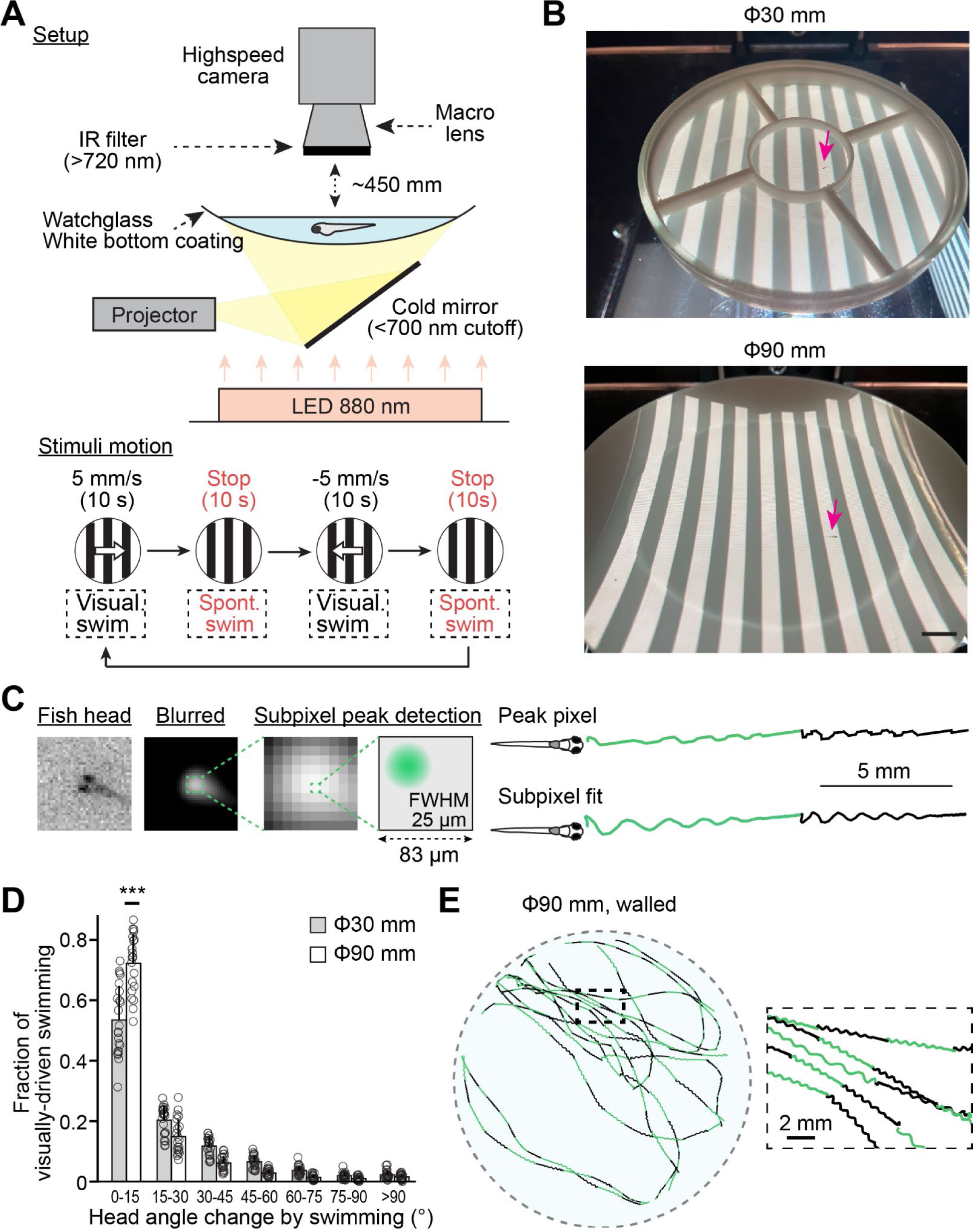
Experimental setup. **(A)** *Top*, our experimental setup for zebrafish tracking in a large environment. Fish behavior is measured in a concave chemical watchglass (for the large environment) or a plastic insert (for the small environment) of which the bottom surface is coated with a white spray. A projector showed visual stimuli in the bottom of the dish at visible wavelength. An infrared illumination was placed under the dish, and a high-speed camera with a macro lens acquire images through an infrared filter. *Bottom*, cycles of visual stimulus motion during the experiment. We define visually-driven swimming as those observed during visual stimulus motion (black) and spontaneous swimming as those observed when the visual stimulus is stopped (red). (**B**) Photos of actual experiments in the small (30 mm) and the large (90 mm) dish, with arrows to help locate the fish. **(C)** Subpixel peak detection significantly improves the accuracy of head centroid trajectories. **(D)** Increased straight swimming during visually-induced swimming in the large arena compared to the small arena. ***, p=1.8*10^-6^ from 2-sample t-test. Error bars represent standard deviations across tested fish. **(E)** Fish’s swim patterns in a large environment (90 mm) with a flat floor and a boundary wall. *Left,* head centroid trajectories of a single fish. *Right,* expanded head centroid trajectories from the outlined central parts of the large arena on the left.

**Supplementary Figure 2:**
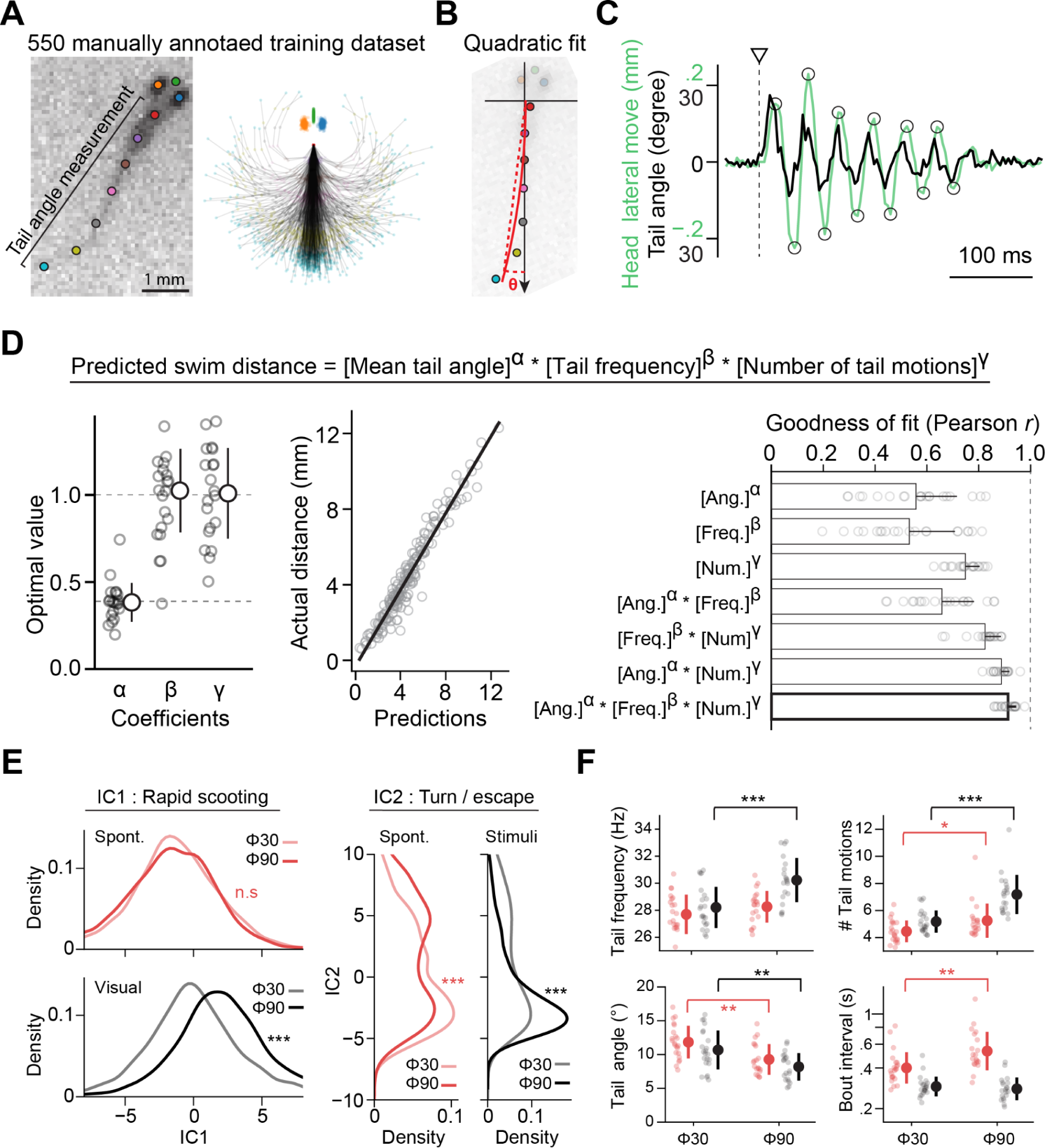
Recognition of tail motions and its accuracy validation. **(A)** *Left*, annotation of 10 points across the body. *Right*, distribution of body parts in 550 manually annotated training datasets. Manual annotations were made using fish images from various orientations, and we aligned them for the visualization of this panel. Training images were selected to balance various tail angles on the left and right sides. **(B)** Tail angle *θ* was quantified by fitting a quadratic function to annotated points along the tail. **(C)** Overlay of tail motions and head centroid motions during a representative swim episode. Peaks of head centroid motions were detected (circles) and used as a reference to quantify tail motions. **(D)** Validating accuracies of tail angle quantification by predicting swimming distance based on tail motion parameters. *Left*, parameters of a multiplicative prediction model (left) were optimized by using an optimizer. *Center*, the resulting model shows high correlations to swimming distance. *Right*, quantification of the prediction accuracy tested in a large environment. The full model (bottom) has an accuracy of Pearson correlation coefficient *r* = 0.89 ± 0.036 across 20 fish. Error bars represent standard deviations. **(E)** Statistical analyses of IC1 and IC2 were performed using the same set of fish in Figures 1 and 2 by using kernel density 2-sample test (see Methods). We included 2,635 (small arena) and 5,233 (large arena) swim episodes for the statistics of spontaneous exploration and 1,466 (small arena) and 5,360 (large arena) swim episodes for the statistics of visually-driven swimming. IC1: n.s. (not significant), p=0.14; ***, p=6.0*10^-49^. IC2: ***, p=1.4*10^-12^ and 1.1*10^-4^ for spontaneous and visually-driven swimming, respectively. **(F)** Analyses of individual swim parameters. The large arena facilitated significantly higher frequencies, more numbers of tail motions and smaller tail angles during visually-driven swimming, while it elongated bout intervals during spontaneous exploration. P values are from a 2-sample t-test between N=22 and N=20 fish for the small and large dishes, respectively. ***, p=2.3*10^-4^ (frequency); *, p=0.021, ***, p=2.8*10^-6^ (motions); ** for spont., p=1.4*10^-3^, ** for visual, p=3.2*10^-3^ (angle); **, p=4.1*10^-3^ (interval). Error bars represent standard deviations across tested fish.

**Supplementary Figure 3:**
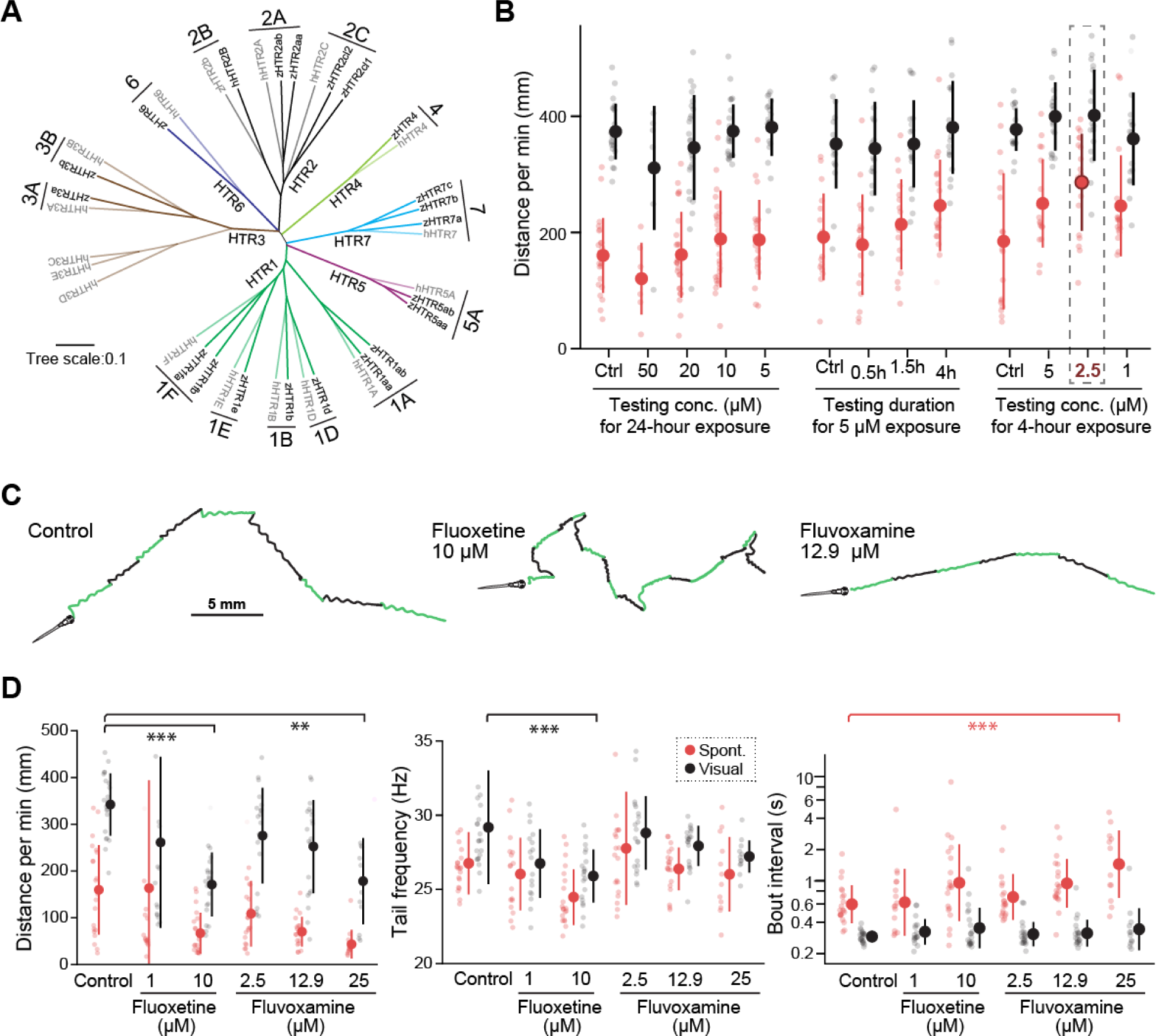
Serotonin receptor families in zebrafish, dose optimization for psilocybin treatment, and behavioral effects of SSRIs. **(A)** Unbiased homology analyses of protein sequences of all serotonin receptors revealed conserved major types and subtypes between zebrafish and humans. **(B)** The effect of psilocybin treatment on swimming distances during spontaneous exploration (red) and visually-driven swimming (black). We tested various durations and concentrations (conc.) of psilocybin exposure. Data from the same set of experimental batches are clustered together with their individual control data. Numbers of fish (left to right): N=22 (Ctrl), 6 (50 μM), 25 (20 μM), 26 (10 μM), 21 (5 μM), 17 (Ctrl), 18 (0.5h), 18 (1.5h), 18 (4h), 18 (Ctrl), 18 (5 μM), 18 (2.5 μM) and 18 (1 μM). The data on the right (4-hour exposure) is the same as those shown in Figure 3E. The condition in the dashed box (2.5 μM, 4 h) had the strongest impact on spontaneous exploration and was used for experiments in Figures 3 and 4. **(C)** Swim trajectories of control, fluoxetine-treated, and fluvoxamine-treated fish during visually-driven behaviors. **(D)** Swimming distance per minute (left), tail frequency (center) and average tail angle (right) of control (N=21), fluoxetine-treated (N=17 for 1 μM, N=20 for 10 μM) and fluvoxamine-treated fish (N=20 for 2.5 μM, N=20 for 12.9 μM, N=12 for 25 μM). P values are from Tukey’s post-hoc test after one-way ANOVA analysis. ***, p=3.8*10^-5^; **, p=1.1*10^-3^ (distance per minute during visually driven swimming); ***, p=6.4 * 10^-4^ (tail frequency during visually driven swimming); ***, p=2.5*10^-4^ (bout intervals during spontaneous exploration). Error bars represent standard deviations.

**Supplementary Figure 4:**
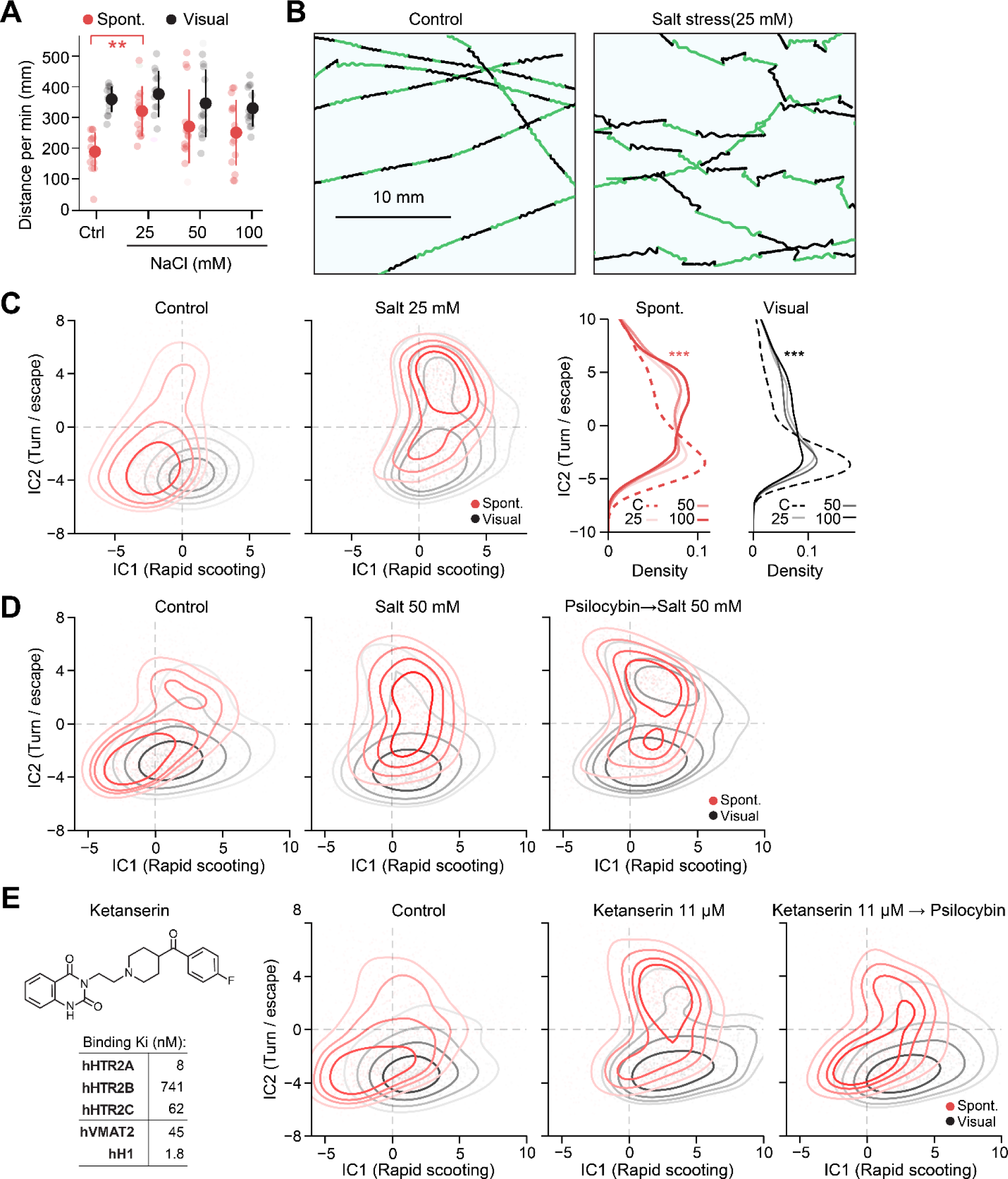
Behavioral changes after exposure to salt stress and ketanserin. **(A)** Swimming distance during spontaneous exploration and visually-driven swimming after exposure to varying concentrations (25 mM, 50 mM and 100 mM) of sodium chloride in the water as hypertonic stress. N=14 fish for all conditions. We used Tukey’s post-hoc test following one-way ANOVA analysis of spontaneous exploration among different conditions for statistics. **, p=0.049. Error bars represent standard deviations. **(B)** Head centroid trajectories around the central part of the large arena after sham treatment (left) and hypertonic stress exposure (right) during visually-driven swimming. **(C)** *Left*, independent component analysis (ICA) revealed the shift of swim patterns toward turn/escape behavior (IC2) after hypertonic stress exposure. The same number (1200) of randomly selected swim events were plotted for each condition. *Right,* statistical analyses of the occurrences of turning/escape behaviors based on IC2 component. We used kernel density 2-sample test (see Methods) for statistics. We included 5,227 (control), 6,259 (25 mM), 5,514 (50 mM) and 5,046 (100 mM) swim episodes for the statistics of spontaneous exploration, and 4,730 (control), 3,322 (25 mM), 3,370 (50 mM) and 3,267 (100 mM) swim episodes for the statistics of visually-driven behavior. ***, p=4.2*10^-28^ (control vs 100 mM for spontaneous exploration); ***, p=4.9*10^-5^ (control vs 100 mM for visually-driven swimming). **(D)** Independent component analysis (ICA) of swim patterns after the sham treatment (left), hypertonic stress exposure (center) and psilocybin pretreatment and stress exposure (right). Data from 11 fish for each condition. **(E)** *Left*, the structure of ketanserin and its affinities to binding targets. Affinity values for HTR2 receptors are from ref^76^, and those for vesicular monoamine transporter 2 (VMAT2) and type 1 histamine receptor (HT1) are from ref^77^ and ref^78^, respectively. *Right*, independent component analysis (ICA) of swim patterns after the sham treatment, ketanserin exposure and psilocybin pretreatment before ketanserin exposure. Data from 14, 15 and 15 fish for each condition, respectively.

## Bibliography

1. Depression. https://www.who.int/news-room/fact-sheets/detail/depression.

2. Coppen, A. The Biochemistry of Affective Disorders. Br. J. Psychiatry 113, 1237–1264 (1967).

3. Dahlstroem, A. & Fuxe, K. Evidence for the Existence of Monoamine-Containing Neurons in the Central Nervous System. I. Demonstration of Monoamines in the Cell Bodies of Brain Stem Neurons. Acta Physiol. Scand. Suppl. SUPPL 232:1–55 (1964).

4. Blakely, R. D. et al. Cloning and expression of a functional serotonin transporter from rat brain. Nature 354, 66–70 (1991).

5. Uher, R. et al. Early and Delayed Onset of Response to Antidepressants in Individual Trajectories of Change During Treatment of Major Depression: A Secondary Analysis of Data From the Genome-Based Therapeutic Drugs for Depression (GENDEP) Study. J. Clin. Psychiatry 72, 5274 (2011).

6. Gaynes, B. N. et al. What Did STAR*D Teach Us? Results From a Large-Scale, Practical, Clinical Trial for Patients With Depression. Psychiatr. Serv. 60, 1439–1445 (2009).

7. Rucker, J. J. H., Iliff, J. & Nutt, D. J. Psychiatry & the psychedelic drugs. Past, present & future. Neuropharmacology 142, 200–218 (2018).

8. Carhart-Harris, R. L. et al. Psilocybin with psychological support for treatment-resistant depression: an open-label feasibility study. Lancet Psychiatry 3, 619–627 (2016).

9. Goodwin, G. M. et al. Single-Dose Psilocybin for a Treatment-Resistant Episode of Major Depression. N. Engl. J. Med. 387, 1637–1648 (2022).

10. Gukasyan, N. et al. Efficacy and safety of psilocybin-assisted treatment for major depressive disorder: Prospective 12-month follow-up. J. Psychopharmacol. (Oxf.) 36, 151–158 (2022).

11. Ross, S. et al. Rapid and sustained symptom reduction following psilocybin treatment for anxiety and depression in patients with life-threatening cancer: a randomized controlled trial. J. Psychopharmacol. (Oxf.) 30, 1165–1180 (2016).

12. Griffiths, R. R. et al. Psilocybin produces substantial and sustained decreases in depression and anxiety in patients with life-threatening cancer: A randomized double-blind trial. J. Psychopharmacol. (Oxf.) 30, 1181–1197 (2016).

13. Daniel, J. & Haberman, M. Clinical potential of psilocybin as a treatment for mental health conditions. Ment. Health Clin. 7, 24–28 (2017).

14. Agin-Liebes, G. I. et al. Long-term follow-up of psilocybin-assisted psychotherapy for psychiatric and existential distress in patients with life-threatening cancer. J. Psychopharmacol. (Oxf.) 34, 155–166 (2020).

15. Daws, R. E. et al. Increased global integration in the brain after psilocybin therapy for depression. Nat. Med. 28, 844–851 (2022).

16. Hibicke, M., Landry, A. N., Kramer, H. M., Talman, Z. K. & Nichols, C. D. Psychedelics, but Not Ketamine, Produce Persistent Antidepressant-like Effects in a Rodent Experimental System for the Study of Depression. ACS Chem. Neurosci. 11, 864–871 (2020).

17. Carhart-Harris, R. L. et al. Functional Connectivity Measures After Psilocybin Inform a Novel Hypothesis of Early Psychosis. Schizophr. Bull. 39, 1343–1351 (2013).

18. Smigielski, L., Scheidegger, M., Kometer, M. & Vollenweider, F. X. Psilocybin-assisted mindfulness training modulates self-consciousness and brain default mode network connectivity with lasting effects. NeuroImage 196, 207–215 (2019).

19. Carhart-Harris, R. L. et al. Psilocybin for treatment-resistant depression: fMRI-measured brain mechanisms. Sci. Rep. 7, 13187 (2017).

20. Shao, L.-X. et al. Psilocybin induces rapid and persistent growth of dendritic spines in frontal cortex in vivo. Neuron 109, 2535–2544.e4 (2021).

21. Cameron, L. P. et al. A non-hallucinogenic psychedelic analogue with therapeutic potential. Nature 589, 474–479 (2021).

22. Dong, C. et al. Psychedelic-inspired drug discovery using an engineered biosensor. Cell 184, 2779–2792.e18 (2021).

23. Cao, D. et al. Structure-based discovery of nonhallucinogenic psychedelic analogs. Science 375, 403–411 (2022).

24. Depping, M. S., Schmitgen, M. M., Kubera, K. M. & Wolf, R. C. Cerebellar Contributions to Major Depression. Front. Psychiatry 9, 634 (2018).

25. Newstead, S. et al. Acute and repetitive fronto-cerebellar tDCS stimulation improves mood in non-depressed participants. Exp. Brain Res. 236, 83–97 (2018).

26. Gouzoulis-Mayfrank, E. et al. Neurometabolic Effects of Psilocybin, 3,4-Methylenedioxyethylamphetamine (MDE) and d-Methamphetamine in Healthy Volunteers A Double-Blind, Placebo-Controlled PET Study with [18F]FDG. Neuropsychopharmacology 20, 565–581 (1999).

27. Ahrens, M. B., Orger, M. B., Robson, D. N., Li, J. M. & Keller, P. J. Whole-brain functional imaging at cellular resolution using light-sheet microscopy. Nat Methods 10, 413–420 (2013).

28. Ahrens, M. B. et al. Brain-wide neuronal dynamics during motor adaptation in zebrafish. Nature 485, 471–7 (2012).

29. Kim, D. H. et al. Pan-neuronal calcium imaging with cellular resolution in freely swimming zebrafish. Nat. Methods 14, 1107–1114 (2017).

30. Lin, Q. et al. Cerebellar Neurodynamics Predict Decision Timing and Outcome on the Single-Trial Level In Brief Article Cerebellar Neurodynamics Predict Decision Timing and Outcome on the Single-Trial Level. Cell 180, 536–551.e17 (2019).

31. Wolf, S. et al. Whole-brain functional imaging with two-photon light-sheet microscopy. Nat. Methods 2015 125 12, 379–380 (2015).

32. Gaspar, P. & Lillesaar, C. Probing the diversity of serotonin neurons. Philos. Trans. R. Soc. B Biol. Sci. 367, 2382–2394 (2012).

33. Parker, M. O., Brock, A. J., Walton, R. T. & Brennan, C. H. The role of zebrafish (Danio rerio) in dissecting the genetics and neural circuits of executive function. Front. Neural Circuits 7, 63 (2013).

34. Kawashima, T. The role of the serotonergic system in motor control. Neurosci. Res. 129, 32–39 (2018).

35. Marques, J. C., Li, M., Schaak, D., Robson, D. N. & Li, J. M. Internal state dynamics shape brainwide activity and foraging behaviour. Nature 577, 239–243 (2020).

36. Kawashima, T. et al. The Serotonergic System Tracks the Outcomes of Actions to Mediate Short-Term Motor Learning. Cell 167, 933–946.e20 (2016).

37. Yokogawa, T., Hannan, M. C. & Burgess, H. a. The dorsal raphe modulates sensory responsiveness during arousal in zebrafish. J. Neurosci. Off. J. Soc. Neurosci. 32, 15205–15 (2012).

38. Filosa, A., Barker, A. J., Dal Maschio, M. & Baier, H. Feeding State Modulates Behavioral Choice and Processing of Prey Stimuli in the Zebrafish Tectum. Neuron 90, 596–608 (2016).

39. Lovett-Barron, M. et al. Ancestral Circuits for the Coordinated Modulation of Brain State. Cell 171, 1411–1423.e17 (2017).

40. Wee, C. L. et al. A bidirectional network for appetite control in larval zebrafish. eLife 8, (2019).

41. Clark, K. J., Boczek, N. J. & Ekker, S. C. Stressing zebrafish for behavioral genetics. Rev. Neurosci. 22, 49–62 (2011).

42. Burgess, H. A. & Granato, M. The neurogenetic frontier—lessons from misbehaving zebrafish. Brief. Funct. Genomics 7, 474–482 (2008).

43. De Marco, R. J., Groneberg, A. H., Yeh, C.-M., TreviÃ±o, M. & Ryu, S. The behavior of larval zebrafish reveals stressor-mediated anorexia during early vertebrate development. Front. Behav. Neurosci. 8, 1–12 (2014).

44. Tan, J. X. M., Ang, R. J. W. & Wee, C. L. Larval Zebrafish as a Model for Mechanistic Discovery in Mental Health. Front. Mol. Neurosci. 15, (2022).

45. Burgess, H. A. & Granato, M. Sensorimotor Gating in Larval Zebrafish. J. Neurosci. 27, 4984–4994 (2007).

46. Andalman, A. S. et al. Neuronal Dynamics Regulating Brain and Behavioral State Transitions. Cell 177, 970–985.e20 (2019).

47. Félix, L. M., Antunes, L. M., Coimbra, A. M. & Valentim, A. M. Behavioral alterations of zebrafish larvae after early embryonic exposure to ketamine. Psychopharmacology (Berl.) 234, 549–558 (2017).

48. Jouary, A., Haudrechy, M., Candelier, R. & Sumbre, G. A 2D virtual reality system for visual goal-driven navigation in zebrafish larvae. Sci. Rep. 6, 34015 (2016).

49. Marques, J. C., Lackner, S., Félix, R. & Orger, M. B. Structure of the Zebrafish Locomotor Repertoire Revealed with Unsupervised Behavioral Clustering. Curr. Biol. CB 28, 181–195.e5 (2018).

50. Severi, K. E. et al. Neural Control and Modulation of Swimming Speed in the Larval Zebrafish. Neuron 83, 692–707 (2014).

51. Zaki, H., Lushi, E. & Severi, K. E. Larval Zebrafish Exhibit Collective Circulation in Confined Spaces. Front. Phys. 9, (2021).

52. Tan, M. & Le, Q. V. EfficientNet: Rethinking Model Scaling for Convolutional Neural Networks. Preprint at https://doi.org/10.48550/arXiv.1905.11946 (2020).

53. Mathis, A. et al. DeepLabCut: markerless pose estimation of user-defined body parts with deep learning. Nat. Neurosci. 21, 1281–1289 (2018).

54. Bialek, W. On the dimensionality of behavior. Proc. Natl. Acad. Sci. 119, e2021860119 (2022).

55. Lenz, C. et al. Assessment of Bioactivity-Modulating Pseudo-Ring Formation in Psilocin and Related Tryptamines. ChemBioChem 23, e202200183 (2022).

56. Glatfelter, G. C. et al. Structure–Activity Relationships for Psilocybin, Baeocystin, Aeruginascin, and Related Analogues to Produce Pharmacological Effects in Mice. ACS Pharmacol. Transl. Sci. 5, 1181–1196 (2022).

57. Erkizia-Santamaría, I., Alles-Pascual, R., Horrillo, I., Meana, J. J. & Ortega, J. E. Serotonin 5-HT2A, 5-HT2c and 5-HT1A receptor involvement in the acute effects of psilocybin in mice. In vitro pharmacological profile and modulation of thermoregulation and head-twich response. Biomed. Pharmacother. 154, 113612 (2022).

58. Halberstadt, A. L. & Geyer, M. A. Multiple receptors contribute to the behavioral effects of indoleamine hallucinogens. Neuropharmacology 61, 364–381 (2011).

59. Nicholas, C. R. et al. High dose psilocybin is associated with positive subjective effects in healthy volunteers. J. Psychopharmacol. (Oxf.) 32, 770–778 (2018).

60. Kirla, K. T. et al. Zebrafish Larvae Are Insensitive to Stimulation by Cocaine: Importance of Exposure Route and Toxicokinetics. Toxicol. Sci. 154, 183–193 (2016).

61. Vollenweider, F. X., Csomor, P. A., Knappe, B., Geyer, M. A. & Quednow, B. B. The Effects of the Preferential 5-HT2A Agonist Psilocybin on Prepulse Inhibition of Startle in Healthy Human Volunteers Depend on Interstimulus Interval. Neuropsychopharmacology 32, 1876–1887 (2007).

62. Carlsson, A. Current theories on the mode of action of antidepressant drugs. Adv. Biochem. Psychopharmacol. 39, 213–221 (1984).

63. Gollan, J. K. et al. What Are the Clinical Implications of New Onset or Worsening Anxiety During the First Two Weeks of SSRI Treatment for Depression? Depress. Anxiety 29, 10.1002/da.20917 (2012).

64. Airhart, M. J. et al. Movement disorders and neurochemical changes in zebrafish larvae after bath exposure to fluoxetine (PROZAC). Neurotoxicol. Teratol. 29, 652–664 (2007).

65. Yang, H. et al. Molecular and behavioral responses of zebrafish embryos/larvae after sertraline exposure. Ecotoxicol. Environ. Saf. 208, 111700 (2021).

66. Griffiths, B. B. et al. A zebrafish model of glucocorticoid resistance shows serotonergic modulation of the stress response. Front. Behav. Neurosci. 6, 68 (2012).

67. Mason, N. L. et al. Psilocybin induces acute and persisting alterations in immune status and the stress response in healthy volunteers. 2022.10.31.22281688 Preprint at https://doi.org/10.1101/2022.10.31.22281688 (2022).

68. Vom Berg-Maurer, C. M., Trivedi, C. A., Bollmann, J. H., De Marco, R. J. & Ryu, S. The Severity of Acute Stress Is Represented by Increased Synchronous Activity and Recruitment of Hypothalamic CRH Neurons. J. Neurosci. Off. J. Soc. Neurosci. 36, 3350–62 (2016).

69. Cheng, R.-K., Tan, J. X. M., Chua, K. X., Tan, C. J. X. & Wee, C. L. Osmotic Stress Uncovers Correlations and Dissociations Between Larval Zebrafish Anxiety Endophenotypes. Front. Mol. Neurosci. 15, (2022).

70. Bai, Y., Liu, H., Huang, B., Wagle, M. & Guo, S. Identification of environmental stressors and validation of light preference as a measure of anxiety in larval zebrafish. BMC Neurosci. 17, 63 (2016).

71. Yeh, C.-M., Glöck, M. & Ryu, S. An Optimized Whole-Body Cortisol Quantification Method for Assessing Stress Levels in Larval Zebrafish. PLOS ONE 8, e79406 (2013).

72. Alderman, S. L. & Bernier, N. J. Ontogeny of the corticotropin-releasing factor system in zebrafish. Gen. Comp. Endocrinol. 164, 61–69 (2009).

73. Shainer, I. et al. Agouti-Related Protein 2 Is a New Player in the Teleost Stress Response System. Curr. Biol. 29, 2009–2019.e7 (2019).

74. Amir-Zilberstein, L. et al. Homeodomain Protein Otp and Activity-Dependent Splicing Modulate Neuronal Adaptation to Stress. Neuron 73, 279–291 (2012).

75. Alsop, D. & Vijayan, M. M. Development of the corticosteroid stress axis and receptor expression in zebrafish. Am. J. Physiol.-Regul. Integr. Comp. Physiol. 294, R711–R719 (2008).

76. De Marco, R. J., Thiemann, T., Groneberg, A. H., Herget, U. & Ryu, S. Optogenetically enhanced pituitary corticotroph cell activity post-stress onset causes rapid organizing effects on behaviour. Nat. Commun. 7, 12620 (2016).

77. Knight, A. R. et al. Pharmacological characterisation of the agonist radioligand binding site of 5-HT2A, 5-HT2B and 5-HT2C receptors. Naunyn. Schmiedebergs Arch. Pharmacol. 370, 114–123 (2004).

78. Kilbourn, M. R. & Koeppe, R. A. Classics in Neuroimaging: Radioligands for the Vesicular Monoamine Transporter 2. ACS Chem. Neurosci. 10, 25–29 (2019).

79. Ghoneim, O. M., Legere, J. A., Golbraikh, A., Tropsha, A. & Booth, R. G. Novel ligands for the human histamine H1 receptor: Synthesis, pharmacology, and comparative molecular field analysis studies of 2-dimethylamino-5-(6)-phenyl-1,2,3,4-tetrahydronaphthalenes. Bioorg. Med. Chem. 14, 6640–6658 (2006).

80. Nowicki, M., Tran, S., Muraleetharan, A., Markovic, S. & Gerlai, R. Serotonin antagonists induce anxiolytic and anxiogenic-like behavior in zebrafish in a receptor-subtype dependent manner. Pharmacol. Biochem. Behav. 126, 170–180 (2014).

81. Müller, T. E. et al. Role of the serotonergic system in ethanol-induced aggression and anxiety: A pharmacological approach using the zebrafish model. Eur. Neuropsychopharmacol. 32, 66–76 (2020).

82. Nic Dhonnchadha, B. Á., Bourin, M. & Hascoët, M. Anxiolytic-like effects of 5-HT2 ligands on three mouse models of anxiety. Behav. Brain Res. 140, 203–214 (2003).

83. León, L. A. et al. Behavioral Effects of Systemic, Infralimbic and Prelimbic Injections of a Serotonin 5-HT2A Antagonist in Carioca High- and Low-Conditioned Freezing Rats. Front. Behav. Neurosci. 11, (2017).

84. Pranzatelli, M. R., Galvan, I. & Tailor, P. T. Human Brainstem Serotonin Receptors: Characterization and Implications for Subcortical Myoclonus. Clin. Neuropharmacol. 19, 507–514 (1996).

85. Pranzatelli, M. R., Tailor, P. T. & Razi, P. Brainstem serotonin receptors in the guinea pig: Implications for myoclonus. Neuropharmacology 32, 209–215 (1993).

86. Norton, W. H. J., Folchert, A. & Bally-Cuif, L. Comparative analysis of serotonin receptor (HTR1A/HTR1B families) and transporter (slc6a4a/b) gene expression in the zebrafish brain. J. Comp. Neurol. 511, 521–542 (2008).

87. Geurts, F. J., De Schutter, E. & Timmermans, J. P. Localization of 5-HT2A, 5-HT3, 5-HT5A and 5-HT7 receptor-like immunoreactivity in the rat cerebellum. J. Chem. Neuroanat. 24, 65–74 (2002).

88. Fay, R. & Kubin, L. Pontomedullary distribution of 5-HT2A receptor-like protein in the rat. J. Comp. Neurol. 418, 323–345 (2000).

89. Zhang, C.-Z. et al. 5-HT2A receptor-mediated excitation on cerebellar fastigial nucleus neurons and promotion of motor behaviors in rats. Pflugers Arch. 466, 1259–1271 (2014).

90. Schneider, H. et al. Cloning and expression of a zebrafish 5-HT2C receptor gene. Gene 502, 108–117 (2012).

91. Romano, S. A. et al. Spontaneous Neuronal Network Dynamics Reveal Circuit’s Functional Adaptations for Behavior. Neuron 85, 1070–1085 (2015).

92. Matsui, H., Namikawa, K., Babaryka, A. & Köster, R. W. Functional regionalization of the teleost cerebellum analyzed in vivo. Proc. Natl. Acad. Sci. 111, 11846–11851 (2014).

93. Hoogland, T. M., De Gruijl, J. R., Witter, L., Canto, C. B. & De Zeeuw, C. I. Role of Synchronous Activation of Cerebellar Purkinje Cell Ensembles in Multi-joint Movement Control. Curr. Biol. 25, 1157–1165 (2015).

94. Barbara, R., Nagathihalli Kantharaju, M., Haruvi, R., Harrington, K. & Kawashima, T. PyZebrascope: An Open-Source Platform for Brain-Wide Neural Activity Imaging in Zebrafish. Front. Cell Dev. Biol. 10, (2022).

95. Vollenweider, F. X. & Kometer, M. The neurobiology of psychedelic drugs: implications for the treatment of mood disorders. Nat. Rev. Neurosci. 11, 642–651 (2010).

96. de Veen, B. T. H., Schellekens, A. F. A., Verheij, M. M. M. & Homberg, J. R. Psilocybin for treating substance use disorders? Expert Rev. Neurother. 17, 203–212 (2017).

97. Lo, R. Y. et al. Depression and Clinical Progression in Spinocerebellar Ataxias. Parkinsonism Relat. Disord. 22, 87–92 (2016).

98. Baek, S. J., Park, J. S., Kim, J., Yamamoto, Y. & Tanaka-Yamamoto, K. VTA-projecting cerebellar neurons mediate stress-dependent depression-like behaviors. eLife 11, e72981 (2022).

99. Wagner, G., de la Cruz, F., Köhler, S. & Bär, K.-J. Treatment Associated Changes of Functional Connectivity of Midbrain/Brainstem Nuclei in Major Depressive Disorder. Sci. Rep. 7, 8675 (2017).

100. Portugues, R. & Engert, F. Adaptive locomotor behavior in larval zebrafish. Front. Syst. Neurosci. 5, 72 (2011).

101. Cong, L. et al. Rapid whole brain imaging of neural activity in freely behaving larval zebrafish (Danio rerio). eLife 6, e28158 (2017).

102. Marvin, J. S. et al. A genetically encoded fluorescent sensor for in vivo imaging of GABA. Nat. Methods 16, 763–770 (2019).

103. Feng, J. et al. A Genetically Encoded Fluorescent Sensor for Rapid and Specific In Vivo Detection of Norepinephrine. Neuron 102, 745–761.e8 (2019).

104. Gross, G. G. et al. Recombinant Probes for Visualizing Endogenous Synaptic Proteins in Living Neurons. Neuron 78, 971–985 (2013).

105. Moro, E. et al. Generation and application of signaling pathway reporter lines in zebrafish. Mol. Genet. Genomics 288, 231–242 (2013).

106. Duong, T., Goud, B. & Schauer, K. Closed-form density-based framework for automatic detection of cellular morphology changes. Proc. Natl. Acad. Sci. 109, 8382–8387 (2012).

107. Sievers, F. & Higgins, D. G. Clustal Omega for making accurate alignments of many protein sequences. Protein Sci. Publ. Protein Soc. 27, 135–145 (2018).

108. Letunic, I. & Bork, P. Interactive Tree Of Life (iTOL) v5: an online tool for phylogenetic tree display and annotation. Nucleic Acids Res. 49, W293–W296 (2021).

109. Choi, H. M. T. et al. Third-generation in situ hybridization chain reaction: multiplexed, quantitative, sensitive, versatile, robust. Development 145, (2018).

110. Shainer, I. et al. A single-cell resolution gene expression atlas of the larval zebrafish brain. 2022.02.11.479024 Preprint at https://doi.org/10.1101/2022.02.11.479024 (2022).

111. Yang, E. et al. A brainstem integrator for self-location memory and positional homeostasis in zebrafish. Cell 185, 5011–5027.e20 (2022).

112. Kunst, M. et al. A Cellular-Resolution Atlas of the Larval Zebrafish Brain. Neuron 103, 21–38.e5 (2019).

113. Avants, B. et al. A reproducible evaluation of ANTs similarity metric performance in brain image registration. NeuroImage 54, 2033–2044 (2011).

